# Ribonanza: deep learning of RNA structure through dual crowdsourcing

**DOI:** 10.1101/2024.02.24.581671

**Authors:** Shujun He, Rui Huang, Jill Townley, Rachael C. Kretsch, Thomas G. Karagianes, David B.T. Cox, Hamish Blair, Dmitry Penzar, Valeriy Vyaltsev, Elizaveta Aristova, Arsenii Zinkevich, Artemy Bakulin, Hoyeol Sohn, Daniel Krstevski, Takaaki Fukui, Fumiya Tatematsu, Yusuke Uchida, Donghoon Jang, Jun Seong Lee, Roger Shieh, Tom Ma, Eduard Martynov, Maxim V. Shugaev, Habib S.T. Bukhari, Kazuki Fujikawa, Kazuki Onodera, Christof Henkel, Shlomo Ron, Jonathan Romano, John J. Nicol, Grace P. Nye, Yuan Wu, Christian Choe, Walter Reade, Eterna participants, Rhiju Das

## Abstract

Prediction of RNA structure from sequence remains an unsolved problem, and progress has been slowed by a paucity of experimental data. Here, we present Ribonanza, a dataset of chemical mapping measurements on two million diverse RNA sequences collected through Eterna and other crowdsourced initiatives. Ribonanza measurements enabled solicitation, training, and prospective evaluation of diverse deep neural networks through a Kaggle challenge, followed by distillation into a single, self-contained model called RibonanzaNet. When fine tuned on auxiliary datasets, RibonanzaNet achieves state-of-the-art performance in modeling experimental sequence dropout, RNA hydrolytic degradation, and RNA secondary structure, with implications for modeling RNA tertiary structure.

## Introduction

RNA molecules that form intricate structures are essential for many biological processes and for emergent RNA-based medicines and biotechnologies. Despite advances in the analogous problem of protein structure prediction,^1,2^ the modeling of an RNA molecule’s structures from its sequence remains unsolved. Recent efforts to predict RNA structure have encountered a number of challenges, including a scarcity of experimentally determined 3D coordinates of RNA structures, the general unavailability of deep multiple sequence alignments, lack of rigor in evaluation, and poor generalization of deep learning models of RNA secondary structure.^3–7^ Several researchers have proposed that chemical mapping experiments, which provide nucleotide-resolution measurements sensitive to RNA structure,^8^ might resolve the current data bottleneck in RNA structure modeling.^9–11^ Nevertheless, there have been no blind tests that would allow rigorous evaluation of whether predictive models might be trained from chemical mapping data.

To resolve these issues, we noted that foundational innovations in deep learning across tasks ranging from vision to natural language processing to protein structure modeling have involved crowdsourcing of datasets and/or model development to large collections of humans coordinated through the internet.^1,16–19^ Motivated by these examples, we here present results from an experiment called Ribonanza (**Fig. 1a**), which involved collection of diverse RNA sequences discovered or collated on the Eterna crowdsourcing platform, acquisition of chemical mapping data on these sequences, and then prospective evaluation of machine learning models on the Kaggle crowdsourcing platform. As in a prior, smaller-scale dual crowdsourcing initiative applied to RNA,^20^ notable aspects of Ribonanza are the diversity of data sources and models, which stems from the large number of human contributors recruited to the challenges; rigorous separation of data designers, experimentalists, and modelers; and evaluation on ‘future’ data that were not available to any participants during modeling (timeline in **Fig. 1a**).

**Figure 1.**
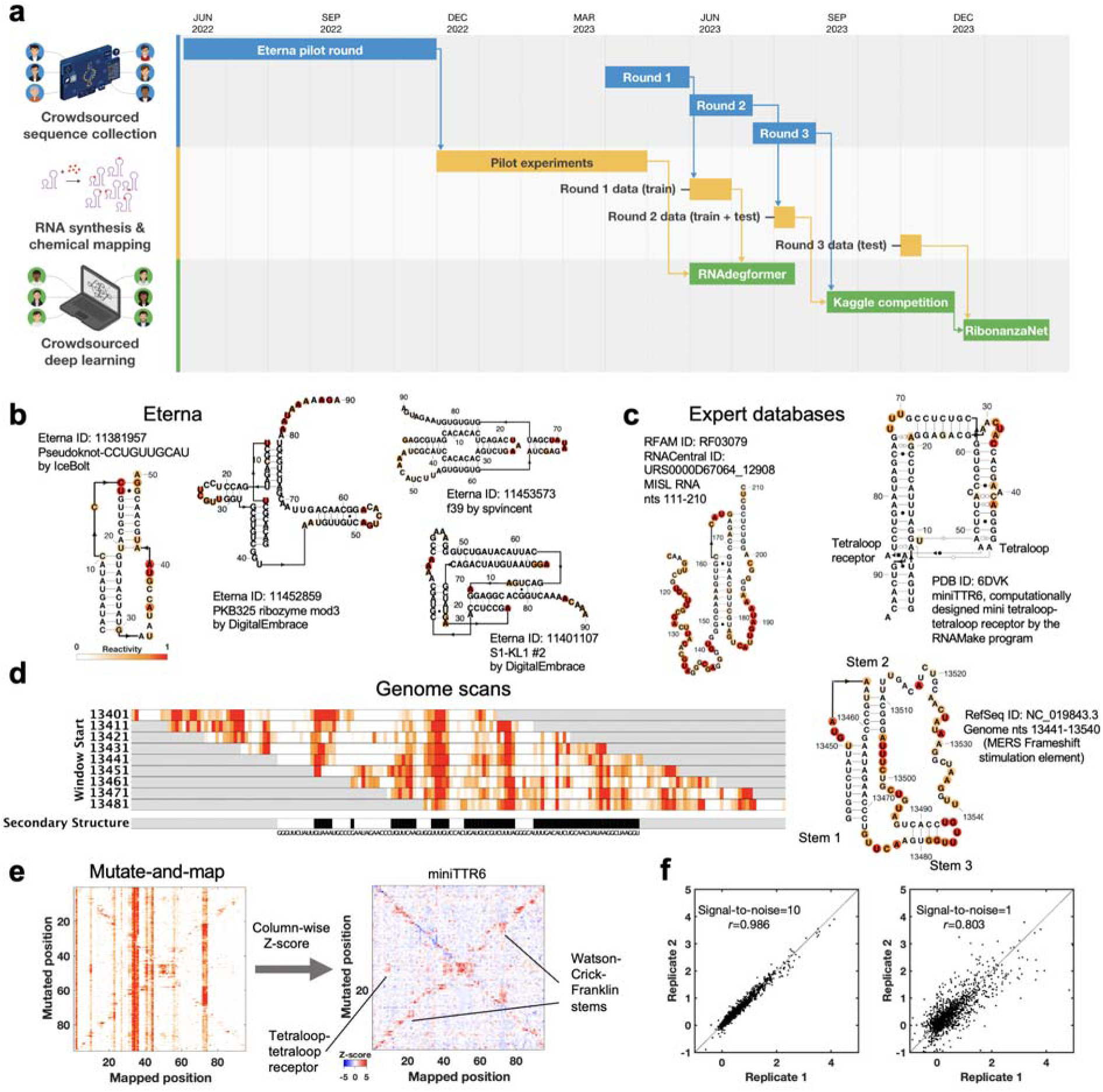
The Ribonanza challenge. **(a)** Timeline of different rounds within the three tracks of Ribonanza: crowdsourced sequence collection, including Eterna design; RNA synthesis and chemical mapping; and crowdsourced deep learning on Kaggle. **(b-c)** Secondary structures with SHAPE (2A3) chemical reactivity data for sequences drawn from **(b)** diverse Eterna submissions and **(c)** expert databases (MISL RNA from RFAM;^12,13^ miniTTR6 nanostructure designed in RNAmake from PDB 6DVK^14^). For each Eterna and RFAM molecule, secondary structures from numerous modeling packages were compared to SHAPE data, and best-fit structure is shown. For miniTTR6, secondary structure and non-canonical base pairs (Leontis-Westhof annotation) were derived from the PDB. **(d)** Reactivity data from a genome scan of Middle Eastern Respiratory Virus; pseudoknotted structure shown on right. **(e)** Mutate-and-map experiments measure reactivity profiles for a sequence mutated at each nucleotide (left); column-wise Z-scores provide more ready visualization of perturbations at sites of mutations as well as at partners involved in Watson-Crick-Franklin stems (secondary structure) and tertiary structure, here shown for miniTTR6. **(f)** Replicate measurements by different experimenters based on DNA template libraries synthesized by different vendors confirm replicability (left); independently measured profiles with estimated mean signal-to-noise ratios as low as 1.0 (right) agree with Pearson’s correlation coefficient *r* > 0.80. Secondary structures in (b)-(d) were prepared in RiboDraw.^15^

## Results

### Diverse RNA molecules with complex predicted structures

As preparatory studies for Ribonanza, we collated sequences from citizen scientists on the Eterna platform^20,21^ (**Extended Data Fig. 1a**) as well as databases curated by experts (e.g., RFAM, the RNA Families database^12^; the PDB archive of 3D coordinates^17^; and Pseudobase^22^); see **Supplemental Tables S2** and **S3**. Current low-cost oligonucleotide synthesis procedures are effective at producing molecules of length 90 to 130 (excluding flanking sequences added to aid experimental characterization; see **Methods**), so longer RNA molecules were segmented into overlapping windows. The largest source of RNA sequences was Eterna, a massive open laboratory that engages a well-established community of citizen scientists in RNA design challenges.^20,21,23^ The Eterna community was recruited through an ‘OpenKnot’ challenge to discover or design RNA sequences with complex structures (**Supplemental Table S2**). The community was given access to computer algorithms that model RNA secondary structures (patterns of Watson-Crick-Franklin base-paired stems). Pseudoknots, which involve pairings of an RNA loop within a stem to partners outside the stem, are hallmarks of complex RNA structure and function^24^ and can be modeled by some of these algorithms (**Methods**). These prior secondary structure modeling methods are known to be only partially accurate^9,25^ and pilot data confirmed the poor predictive power of models, especially deep learning models (**Extended Data Fig. 1b-c**). Nevertheless, we reasoned that acquiring experimental data on sequences predicted to fold into complex structures would provide useful data for refinement of the existing models and training and evaluation of better ones.

The RNA sequences collected through these efforts were synthesized and subjected to chemical mapping methods based on dimethyl sulfate (DMS)^26,27^ and SHAPE (selective 2’ hydroxyl acylation and primer extension) with the modifier 2A3,^28,29^ which mark nucleotides that are not sequestered into base pairs. Pilot studies confirmed that many RNA sequences compiled from different sources, including Eterna, RFAM, and the PDB gave chemical mapping profiles consistent with pseudoknots or non-canonical tertiary contacts and inconsistent with simpler secondary structures (**Fig. 1b-c**, **Extended Data Fig. 1d**, and **Methods**). Genome scans revealed highly structured regions; synthesizing the genome in windows captured known pseudoknots like the frameshift stimulation elements of SARS-CoV-1, MERS, and SARS-CoV-2 coronaviruses which can adopt alternative folds in a larger genomic context (**Fig. 1d**).^30,31^ For some RNA molecules, we also included collections of mutate-and-map (M^2^) sequences.^32^ These experiments monitor how the chemical reactivities at each position in an RNA are affected by mutations introduced at every other position in the RNA, revealing patterns corresponding to secondary structure and, in favorable cases, tertiary contacts (**Fig. 1e**).^33,34^

These pilot studies also confirmed that for libraries with up to 120,000 different RNA sequences, the majority of molecules could be profiled through Illumina sequencing with excellent reproducibility (r^2^>0.8) between independent replicates prepared from libraries synthesized by different commercial providers (**Fig. 1f**). Furthermore, a simple signal-to-noise estimate based on statistical error in Illumina sequencer read counts allows distinction between profiles with acceptable or poor reproducibility and scales in a predictable manner with the number of sequencing reads (**Methods**; **Extended Data Fig. 2**). We chose to further scale up to libraries with up to 1,000,000 sequences, with the expectations that at least 10% of the RNA molecules would still have high signal-to-noise and that noisy data on the remaining sequences might still provide useful signal for machine learning.

### Automatic deep learning of RNA structure representations

RNA structure models have historically been developed and tested through comparisons of model predictions to small sets of hundreds to thousands of secondary structures curated by experts based on available sequence alignments. These datasets have focused on special RNA molecules that have single dominant structures; most RNA molecules however do not have this property. Prior work has demonstrated how a modeling algorithm (EternaFold) can be trained and evaluated via direct comparison to nucleotide-resolution biochemical data on molecules that potentially form multiple structures, though that work was limited to a conventional secondary structure prediction model with strong biophysical assumptions that, e.g., disallowed pseudoknot modeling.^23^ Here we tested whether RNA structure models based on deep learning with few or no prior assumptions might be trained and evaluated based on chemical mapping data. Our first tests came from experiments carried out on a set of 1.1 million RNA sequences described above, including Pilot Round, Round 1, and Round 2 of the Eterna OpenKnot challenge. For model training, we separated out sequences drawn from 17 sublibraries involving Eterna, Rfam, RNAmake designs, and mutate-and-map sequences; another 6 diverse sublibraries provided test sets (**Supplemental Table S4**). Within the training set, 170,000 sequences achieved signal-to-noise greater than 1.0 in both DMS and 2A3 experiments (**Extended Data Fig. 2** and **Supplemental Table S4**). We used data for 140,000 sequences for experiments, reserving the rest as validation data to monitor convergence of model training (**Methods)**.

To understand whether chemical mapping data at this scale might enable training of predictive deep learning models, we used an architecture called RNAdegformer, previously developed for modeling chemical degradation of RNA sequences from in-line hydrolysis in the OpenVaccine effort (**Fig. 2a**).^35^ RNAdegformer involves the Transformer encoder,^36^ whose blocks process a one-dimensional representation of the sequence. In prior work, best predictions from RNAdegformer came from supplementing standard Transformer operations with one dimensional convolutional operations, which are effective in capturing information on sequence-local motifs, and biasing the pairwise attention matrix with terms encoding sequence distance as well as the base pair probability (BPP) matrix computed by conventional secondary structure prediction methods like EternaFold. Additionally, inspired by strategies from the OpenVaccine Kaggle competition,^20,36^ we augmented training data by flipping sequences (reading sequence from 3’-to-5’ and 5’-to-3’) along with their reactivity data. We found that RNAdegformer models could be trained to convergence within hours on widely available graphics processing units, with more training time required for ‘sequence-only’ models that did not use EternaFold BPP information (**Methods** and **Supplemental Table S5**). Ablation studies confirmed that RNAdegformer models trained with lower amounts of data, without the EternaFold BPP matrix, and without flip augmentation gave worse quantitative agreement with test data (**Fig. 2b**).

**Figure 2.**
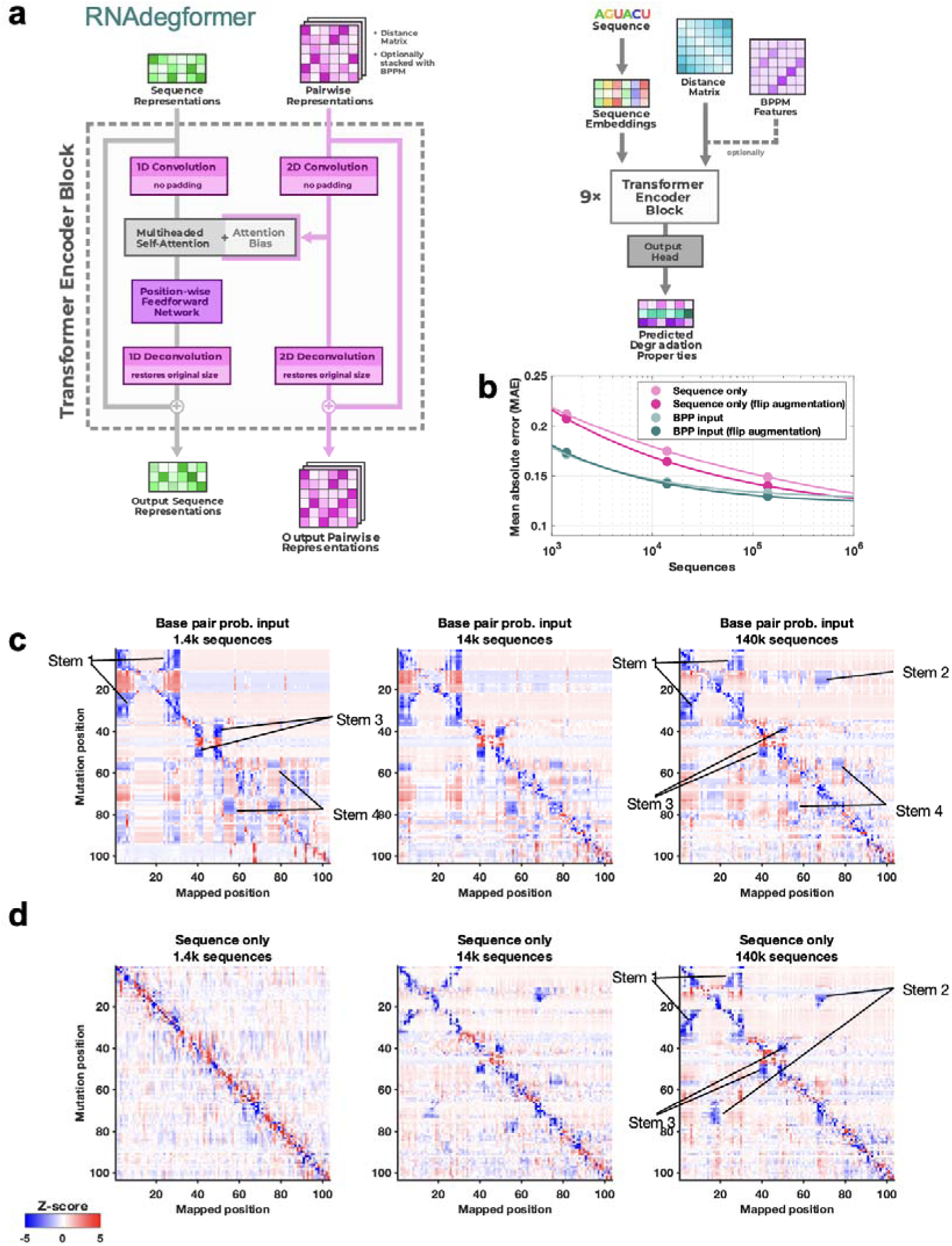
Realistic representations of RNA structure learned from chemical mapping data. **(a)** RNAdegformer model consists of Transformer encoder layers supplemented by convolutions. Attention matrices are biased by sequence-distance matrices and, optionally, base pairing probability (BPP) matrices from conventional secondary structure modeling algorithms, here EternaFold. **(b)** Increasing training data improves RNAdegformer modeling accuracy more rapidly for sequence-only models than for models with BPP input. MAE is mean absolute error on chemical reactivity, after clipping data and predictions to values between 0.0 and 1.0. The curve fits assume a simple power law, MAE = *a* + *b* Sequences^−*c*^, where *a*, *b*, and *c* are fit parameters. **(c-d)** Mutate-and-map predictions for the MERS frameshift stimulation element by RNAdegformer models trained with increasing amount of chemical mapping data either **(c)** with or **(d)** without BPP input.

To evaluate whether an RNAdegformer model might develop internal representations of RNA structure from chemical mapping data, we took advantage of the M^2^ approach, here applied *in silico*. The MERS coronavirus frameshift stimulation element (FSE), has been proposed to form a three-stem pseudoknot secondary structure^30,31,37^ that is consistent with SHAPE data, which were available for the wild type sequence (but not for M^2^ mutants) in our pilot experiments (**Fig. 1d**). RNAdegformer models predicted realistic M^2^ maps in which *in silico* mutations of stems gave rise to reactivity changes in base pairing partners (**Fig. 2c**). When trained with only 1,400 or 14,000 sequences, the features appeared for a non-pseudoknotted structure that misses Stem 2, reflecting the biases of the EternaFold model, which cannot model pseudoknotted secondary structures. When trained with all 140,000 data, however, the M^2^ predictions showed evidence of the expected Stem 2 and a pseudoknotted structure (**Fig. 2c**, middle and right).

Results with the ‘sequence-only’ RNAdegformer, without EternaFold BPP input, were different. When trained on 1,400 sequences this model gave no features representing Watson-Crick-Franklin stems (**Fig. 2d**, left). However, training the model on ten-fold more (14,000) sequences led to a predicted M^2^ map with weak features corresponding to FSE stems (**Fig. 2d**, middle). Training on 140,000 sequences led the RNAdegformer to rebalance these features, with three stems strengthened and the fourth stem largely disappearing, leaving a three-stem pseudoknot (**Fig. 2d**, right). These studies confirmed that models can learn representations of RNA structure, including pseudoknots, motivating prospective tests, including full experimental M^2^ data sets for the MERS FSE, described next.

### Crowdsourcing on Kaggle elicits diverse deep learning models

To advance and rigorously evaluate deep learning models based on chemical mapping data, independent groups were recruited to develop models through a three-month blind competition on the Kaggle platform (**Fig. 3a**). At the beginning of this competition, the data described above on 1.1 million RNA sequences of lengths ranging from 115 to 206 were available. Out of these sequences, 300,000 test molecules were held out as a public test set to allow for continuous evaluation by teams through the competition (‘Public Leaderboard’; see also **Supplemental Table S4**). This evaluation set was additionally filtered to ensure high signal-to-noise and read coverage (**Methods**). Data from 2A3 and DMS mapping for the remaining 800,000 sequences were made available to Kaggle teams for model training. Separate from the training data and continuous evaluation data (Public Leaderboard), a set of 1 million sequences were reserved for final evaluation (Private Leaderboard), seeded with data for 20,000 sequences that were available at the beginning of the competition (**Supplemental Table S4**). To ensure rigor, the vast majority of this private test set was experimentally synthesized and profiled after the Kaggle competition began. Furthermore, to encourage development of models with length generalization, these ‘future’ sequences had lengths ranging from 207 to 457 nucleotides, intentionally chosen to have a different length distribution compared to the training data. A mean absolute error (MAE) metric was chosen for evaluation of models to help reduce impact of outliers, based on tests with submissions collected in the OpenVaccine Kaggle competition (**Extended Data Fig. 3a**). Precomputed EternaFold BPP matrices were made available for all sequences. In addition, predictions of other structure modeling packages as well as a curated dataset of chemical mapping profiles of 70,000 sequences from the RNA Mapping Database were made available (see **Methods**) but did not turn out to be useful for top competitors.

**Figure 3.**
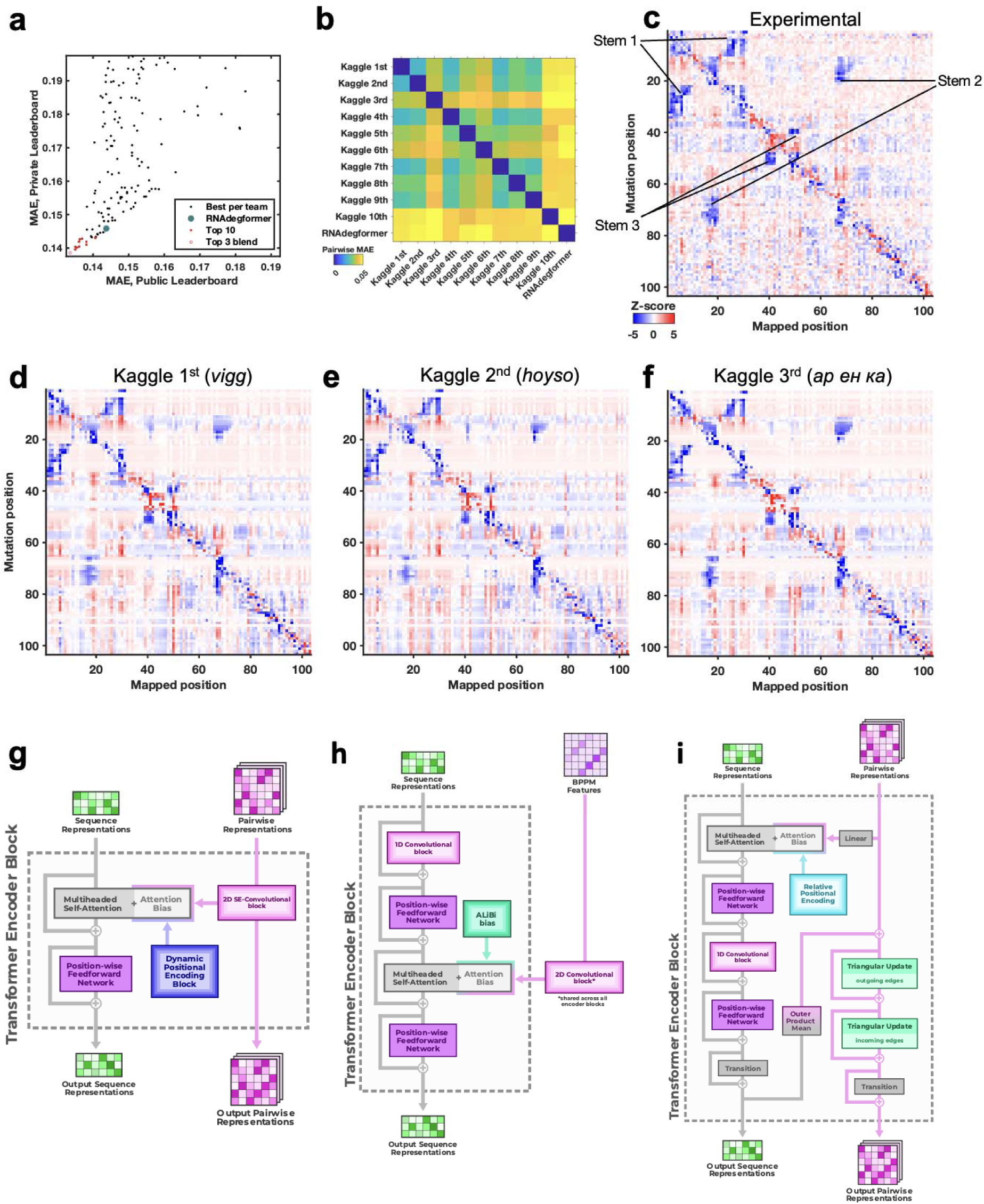
Diverse deep learning models from Kaggle Ribonanza challenge. **(a)** Scores on test sets used for continuous evaluation (Public Leaderboard) vs. prospective evaluation (Private Leaderboard). MAE is mean absolute error compared to experimental chemical reactivity data, after clipping data and predictions between 0.0 and 1.0. **(b)** MAE between different models suggests diversity in predictions even within top 10 Kaggle models. **(c-f)** Mutate-and-map data for the MERS frameshift stimulation element, as measured through SHAPE/2A3 experiments and predicted by 1st, 2nd, and 3rd place Kaggle models. To obtain sufficient signal-to-noise, each row in **(c)** averages over all sequences that harbored a mutation at the corresponding position. **(g-i)** Transformer encoder operations for **(g)** 1st place model (team *vigg*), **(h)** 2nd place model (team *hoyso*), and **(i)** one of two models used for 3rd place submission (*ар ен ка*). Diagrams of full architectures provided in **Extended Data Fig. 6**.

The Kaggle competition recruited 891 participants in 755 teams, and 20 of these teams made predictions that outperformed the RNAdegformer benchmark (**Fig. 3a**). Amongst the top 50 teams, the rankings on the continuous evaluation data (Public Leaderboard) closely matched final rankings (Private Leaderboard) that were based primarily on subsequently collected data from longer sequences (**Fig. 3a**; Spearman *r*_s_ = 0.82), supporting good model generalization and the choice of MAE as an evaluation metric (**Extended Data Fig. 3b**). These teams also produced submissions that were closer to each other (lower MAE) than to the RNAdegformer baseline (**Fig. 3b**). This increased precision suggested that models were able to make more effective use of training data than the baseline model. The top models also appeared to have independently developed realistic representations of RNA structure. **Fig. 3c** shows the experimental M^2^ data acquired for the MERS FSE as part of the test data for the Kaggle challenge; the map clearly shows features for a three-stem pseudoknot, confirming predictions of RNAdegformer (**Fig. 2d**). Blind predictions of top Kaggle models (**Fig. 3d-f**) also produced M^2^ maps that were nearly indistinguishable from each other and from the experimental data. Although giving consistent results on the MERS FSE, M^2^ predictions of different Kaggle models disagreed for other test cases. **Extended Data Fig. 4** compares results for the *Tetrahymena* group I ribozyme in which different pseudoknots were predicted by the Kaggle models, with no model being completely accurate, and a tetraloop/tetraloop-receptor tertiary interaction was not clearly captured in most Kaggle models, despite this interaction being displayed in molecules in the training set (**Figs**. **1c,e**).

Beyond the *Tetrahymena* ribozyme, additional analyses highlighted differences between modeling approaches. For example, while the first place model by team *vigg* performed best across many sublibraries of the test set, the 3rd place model by *ар ен ка* performed better for other sublibraries (**Extended Data Fig. 5**), suggesting that they might have core differences in their approaches. In addition, blending the top 3 submissions with equal weights produced a model with better accuracy than the individual models (open red circle, **Fig. 3a**), suggesting that independent Kaggle teams developed independent innovations.

Descriptions and code for top models released by groups after the competition confirmed similarities but interesting differences between model architectures (**Fig. 3g-i**, **Extended Data Fig. 6**, **Supplemental Table S5**). As is common in Kaggle competitions, the top submissions all involved linear combinations (‘blends’) of multiple predictions, often made by a single model architecture with different parameter settings. **Fig. 3g** shows the main architecture of the first place solution by team *vigg*, which was similar to other top 10 solutions and the RNAdegformer baseline (**Fig. 2a**; **Supplemental Table S5**) in that it accepted input RNA sequences, processed features through attention layers and residue-wise feed-forward neural networks typical of Transformer encoders, and outputted predicted chemical reactivities. Similar to RNAdegformer, this and other top models supplemented Transformer layers with convolutional operations, used relative position encodings, and biased self-attention matrices with pairwise representations (**Supplemental Table S5)**. In setting up the pairwise representations, all of the top submissions used EternaFold BPP features as inputs, though the submission from third place team *ар ен ка* included one model (out of two) that did not use EternaFold BPP. This twin tower model (**Fig. 3i**) was distinct from RNAdegformer and other Kaggle models – but similar to protein modeling networks AlphaFold2^1^ and RoseTTAfold^4^– in that it passed information from a track (‘tower’) with one dimension corresponding to the RNA sequence, to the pair track, which has two such sequence dimensions.

To achieve increased accuracy and accelerate further application of Ribonanza deep learning models, we sought to develop a single, self-contained model that integrated interesting features of the different Kaggle models. It was particularly intriguing that, while most Kaggle submissions utilized EternaFold BPP matrices, one of two models contributing to the third place solution of *ар ен ка* did not use this shortcut. Prompted by this observation, we developed a sequence-only model called RibonanzaNet which would not require use of BPP features but might make better use of 1D representations to update pair features (**Fig. 4a** and **Methods**). We also explored using pseudo labels derived from the top 3 Kaggle predictions to improve RibonanzaNet accuracy (**Methods** and **Extended Data Fig. 7**). The resulting model surpassed the Kaggle first place solution (**Extended Data Fig. 7**). The result was particularly striking since all of the top Kaggle submissions, including the first place solution, involved blending predictions from numerous models, while RibonanzaNet is a single model. Parallel efforts by Kaggle 1st place team *vigg* showed that another single model, called ArmNet (Artificial Reactivity Mapping using neural Networks), led to improved performance upon integration of concepts from prior models and tested the necessity of ‘triangular’ operations explored in RibonanzaNet (**Methods, Supplemental Table S5,** and **Extended Data Fig. 8**).

**Figure 4.**
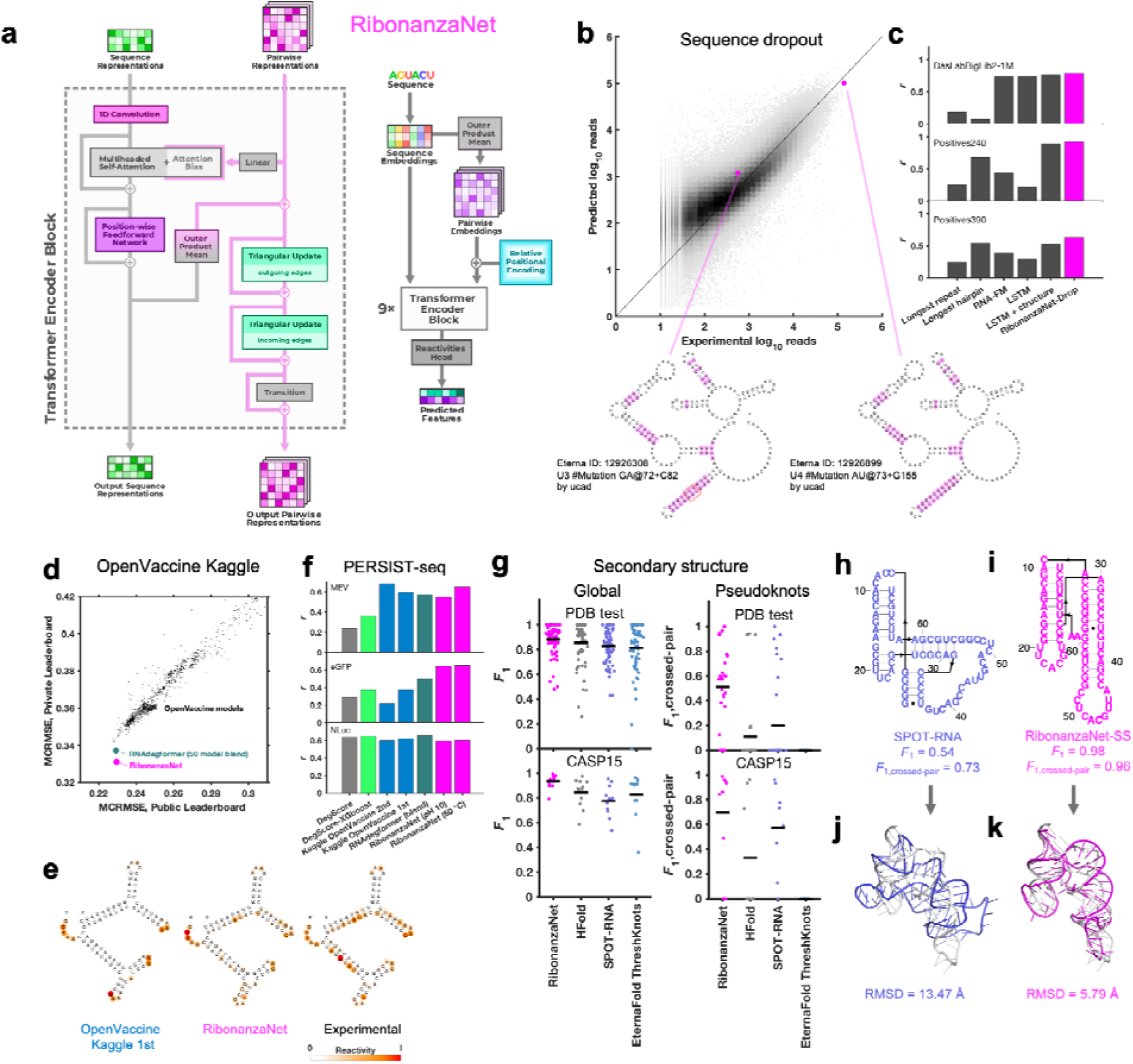
RibonanzaNet model and fine-tuning for downstream tasks. **(a)** RibonanzaNet architecture unifies features of RNAdegformer and top Kaggle models into a single, self-contained model. **(b)** Predictions of RibonanzaNet-Drop for sequence dropout during SHAPE chemical mapping experiments, tested on DasLabBigLib2-1M after fine-tuning on Illumina sequence read counts in DasLabBigLib-1M. Diagrams depict similar sequences (differences highlighted in magenta; red rounded rectangle in left-hand diagram show G-C pairs stabilizing long stem) with identical predicted secondary structures (pseudoknot not shown) but different levels of dropout. **(c)** Pearson correlation coefficients of logarithm of sequencer read counts compared to RibonanzaNet-Drop and baseline models for three test sets. **(d)** Modeling accuracy of RibonanzaNet-Deg for OpenVaccine test sets (Public & Private Leaderboard) after fine-tuning on OpenVaccine training examples. MCRMSE: mean column root mean squared error for SHAPE reactivity and two degradation conditions (pH 10 and 50 °C, 10 mM MgCl_2_, 1 day). **(e)** SHAPE reactivity of OpenVaccine test molecule ‘2204Sept042020’ predicted by top model in Kaggle OpenVaccine competition (‘Nullrecurrent’) and RibonanzaNet, compared to experimental profile. **(f)** Pearson correlation coefficients of degradation profiles predicted by different algorithms compared to degradation rates measured by PERSIST-seq for mRNA molecules encoding a multi-epitope vaccine SARS-CoV-2 (MEV), enhanced green fluorescent protein (eGFP), and nanoluciferase (NLuc). **(g)** Secondary structure accuracies of RibonanzaNet-SS and other packages on a temporally split PDB test set (top) and CASP15 RNA targets (bottom). *F*_1_ is harmonic mean of precision and recall of base pairs; lines in violin plots display mean *F*_1_. **(h-i)** Secondary structure models for CASP15 target R1107, human CPEB3 ribozyme, derived from (**h)** SPOT-RNA and **(i)** RibonanzaNet.^38^ **(j-k)** Overlay of X-ray structure (white) and 3D models using trRosettaRNA guided by secondary structures from **(j)** SPOT-RNA and **(k)** RibonanzaNet.

### Tests of RibonanzaNet on downstream tasks: sequence dropout, RNA degradation, and structure prediction

After achieving better MAE than all solutions from the Ribonanza Kaggle challenge for predicting chemical mapping measurements, we tested whether the RibonanzaNet model might be useful for four distinct tasks: modeling dropout of sequences in our experiments, RNA degradation from hydrolysis, RNA secondary structure, and RNA tertiary structure.

The first task we undertook was development of a quantitative model of the dropout of sequences during chemical mapping experiments. Such a model would enable difficult sequences to be avoided in future experiments, but is expected to depend in a complex manner on structure, sequence repeats, and numerous unknown factors. Prior to the availability of Kaggle models, we explored models based on heuristics involving hairpin lengths and sequence repeats as well as long-short term memory networks^39^ and the foundation model RNA-FM^40^, trained on Illumina sequencer read counts for a library of 1 million 177-nt sequences available at the beginning of the competition (**Methods)**. When fine-tuned to the same training set, RibonanzaNet-Drop gave better correlation coefficients to experimental logarithm of read counts than any of the baseline models on three test sets with molecules of longer lengths (**Fig. 4b-c** and **Supplemental Table S6**). Interestingly, test molecules with similar sequence and identical predicted secondary structures gave strikingly different levels of dropout experimentally. These differences were accounted for by the RibonanzaNet-Drop model and appear to be caused by increased stability of G-C base pairs in the poorly represented sequence, which would decrease efficiency of reverse transcription (**Fig. 4b**, bottom).

Second, we turned our attention to OpenVaccine, the previous Eterna/Kaggle dual crowdsourcing effort, which sought models of RNA hydrolytic degradation with the goal of aiding design of more thermostable mRNA vaccines.^20^ For this application, we prepared RibonanzaNet-Deg, fine-tuned to predict SHAPE and degradation profiles using the same training dataset used in OpenVaccine (**Methods**). On the test dataset, RibonanzaNet-Deg achieves markedly lower MCRMSE (mean column root mean squared error, the evaluation metric used in OpenVaccine) for the OpenVaccine competition’s private leaderboard test set, compared to the top OpenVaccine solutions as well an RNAdegformer model ensemble developed after OpenVaccine (**Fig. 4d**).^35^ **Fig. 4e** shows an example test molecule ‘2204Sept042020’ that was a problem case previously; unlike top OpenVaccine models, RibonanzaNet-Deg models the molecule’s A/U-rich stems as unfolded and degradation-prone, in agreement with experiment. As a further test, RibonanzaNet outperforms the prior models in recovering degradation rates of completely different sets of longer mRNA molecules, measured through PERSIST-seq (**Fig. 4f**).^20,41^

Third, we pursued prediction of RNA secondary structure, particularly the pseudoknot structures that are common in functional RNA. Accurate modeling of the Watson-Crick-Franklin base pairing of nucleotides is a prerequisite for most current 3D structure prediction methods but has required human intervention in blind competitions like the recent CASP15 experiment.^5,9,25,42–44^ To address this challenge, we fine-tuned RibonanzaNet on secondary structures extracted from PDB coordinates of experimentally resolved tertiary structures available prior to May 2020. We tested this RibonanzaNet-SS model on subsequent PDB depositions as well as CASP15 RNA targets using the *F*_1_ metric, the harmonic mean of base pair precision and recall (**Methods**). On the PDB test dataset, RibonanzaNet-SS outperforms previous state-of-the-art algorithms (mean *F*_1_ of 0.890 vs. 0.867 on next best algorithm, HFold from Shapify, **Fig. 4g** and **Supplemental Table S7**; comparisons were only made among algorithms trained without multiple sequence alignments). Fine tuning the model without pre-training gave worse results (mean *F*_1_ 0.710; **Supplemental Table S7**), confirming the importance of Ribonanza data to enable successful transfer learning. On twelve CASP15 RNA-only secondary structures,^9^ RibonanzaNet-SS achieves a particularly high average *F*_1_ of 0.936 (**Fig. 4g, Supplemental Tables S7, S8**).

To more specifically evaluate complex secondary structures, we developed a metric that focuses on pseudoknots, which are stems whose loops pair to nucleotides outside the stem (i.e., base pairs that ‘cross’ other base pairs; see **Methods**). By this average *F*_1, crossed-pair_ measure, RibonanzaNet-SS achieves higher performance compared to the next best pseudoknot predictor, SPOT-RNA,^45^ on both the PDB test dataset (0.496 vs. 0.272) and the CASP test set (0.698 vs. 0.590) **(Fig. 4g** and **Supplemental Table S7**). Analogous to confidence measures in wide use for 3D modeling, we noted that a simple combination of RibonanzaNet pair scores allowed for accurate estimates of expected modeling accuracy for secondary structure over all base pairs and just pseudoknotted pairs, called *eF*_1_ and *eF*_1,crossed-pair_, respectively (**Methods** and **Extended Data Fig. 9**). These scores allow for flagging of both non-confident and confident predictions (e.g., for the *Tetrahymena* ribozyme and MERS FSE, respectively; **Extended Data Fig. 4h-i** and **Extended Data Fig. 10**). Data ablation studies, including use of a higher quality Ribonanza+ data set acquired after the Kaggle competition, suggest that further increases in chemical mapping data should lead to continuing increases in average *F*_1, crossed-pair_ (**Extended Data Fig. 11**).

Finally, we investigated improving RNA tertiary structure prediction. The top deep learning model in CASP15 based on root mean squared deviation (RMSD) 3D accuracy, trRosettaRNA, used secondary structure predictions from SPOT-RNA that were inaccurate for several targets.^3^ **Fig. 4h-i** shows secondary structures of the CASP15 target R1107, the human CPEB3 ribozyme, for which RibonanzaNet-SS predictions achieve nearly double the accuracy of the deep learning algorithm SPOT-RNA.^45^ We compared the accuracy of trRosettaRNA 3D predictions when utilizing BPP matrices from either SPOT-RNA (the default method utilized by trRosettaRNA) or RibonanzaNet-SS. With RibonanzaNet-SS information, the RMSD to experimental 3D coordinates improved dramatically in some targets, including R1107 (**Fig. 4j-k**). However, RMSD improvements were not consistent across all test molecules (**Extended Data Fig. 12, Supplemental Table S8**). We noted that 3D modeling with trRosetaRNA generally deteriorated accurate secondary structures from RibonanzaNet-SS (e.g., average *F*_1, crossed-pair_ dropped from 0.496 to 0.234 on the PDB test dataset). Alternative 3D folding methods, like RoseTTaFold and AlphaFold3,^6,7^ may better leverage information from RibonanzaNet but currently do not accept RNA secondary structure as input.

## Discussion

Motivated by data bottlenecks and slow progress in RNA structure modeling, Ribonanza is an internet-scale, open science project involving the collection of chemical mapping measurements integrated with the development and prospective assessment of deep learning models trained and tested on these data. Dual crowdsourcing through the Eterna and Kaggle platforms enabled inclusion of diverse natural and synthetic designs and diverse machine learning architectures, respectively. Top Kaggle models outperformed previously available models on test data collected after the start of the Kaggle competition. In tests based on mutate-and-map experiments, these models show evidence of learning representations of RNA secondary structure, including pseudoknots. Integrating compelling features of different models into a single RibonanzaNet architecture produces better predictions than the separate models and obviates the need for base pairing probability features derived from conventional RNA secondary structure modeling algorithms. RibonanzaNet provides predictions for chemical mapping data that can be immediately used for guiding RNA design and discovery of novel RNA structures, and fine-tuned networks appear accurate in tasks as disparate as identifying sequences that drop out of experiments, predicting RNA hydrolytic degradation, and inferring RNA secondary structure for 3D modeling.

There are important limitations to our study. We were not able to prospectively assess performance on 3D structure prediction due to the time and expense required to train neural 3D structure modules and to solve novel structures by crystallography, NMR, or cryo-EM. Such assessment awaits the completion of the ongoing CASP16 competition and future RNA-Puzzles trials. In addition, the current datasets involve approximately 400 million nucleotide-level measurements for two million sequences, with acceptable signal-to-noise achieved for 80 million measurements. These datasets are much smaller than the corpora of 10 billion to many trillions of data that have led to transformative deep learning models in areas like natural language processing.^46^ Increased scaling of chemical mapping measurements is technically feasible and is expected to lead to more accurate deep learning models, but remains to be demonstrated.

## Author contributions

J.T., J.R., T.G.K., and R.D. designed and coordinated the Eterna OpenKnot challenge, and Eterna participants contributed sequences through this challenge. R.C.K., J.J.N., and R.D. developed and applied software for collating Ribonanza sequence libraries; J.T., R.C.K., G.P.N., C.C., and R.D. collated additional sequences for Ribonanza. R.H. and R.D. developed, coordinated, and analyzed chemical mapping experiments; and R.H. and Y.W. executed experiments. S.H., W.R., and R.D. designed the Ribonanza Kaggle competition. S.H., D.P., V.V., E.A., A.Z., A.B., H.S., D.K., T.F., F.T., Y.U., D.J., J.S.L., R.S., T.M., E.M., M.V.S., H.S.T.B., K.F., K.O., C.H., and S.R. developed models in the Kaggle competition, and S.H., J.R., and R.D. analyzed submissions. D.P., V.V., E.A., A.Z., and A.B. developed ArmNet. S.H. developed and trained RibonanzaNet and RibonanzaNet-Deg; S.H. and D.B.T.C. developed RibonanzaNet-SS; H.M.B. developed RibonanzaNet-Drop; and D.B.T.C. and S.H. performed 3D modeling studies. R.D. coordinated writing of the manuscript by all authors.

## Acknowledgments

We thank M. Demkin and A. Chow (Kaggle) for expert administration of the Kaggle competition; J. Yesselman and C. Geary for aid in designing 3D nanostructures; D. Incarnato, and A. Mustoe for sharing chemical mapping insights and code; and A.M. Watkins and R. Wellington-Oguri for advice on Eterna puzzles. This work was funded by a Texas A&M University X-grant (S.H.), the National Institutes of Health (R01 AI165433 to S.H; R35 GM122579 to R.D.), and Howard Hughes Medical Institute (to R.D.). This article is subject to HHMI’s Open Access to Publications policy. HHMI lab heads have previously granted a nonexclusive CC BY 4.0 license to the public and a sublicensable license to HHMI in their research articles. Pursuant to those licenses, the author-accepted manuscript of this article can be made freely available under a CC BY 4.0 license immediately upon publication.

## Methods

### Eterna OpenKnot challenge

The design of RNA molecules in Eterna through on-line ‘puzzles’ has been described previously.^15^ Pilot rounds of Eterna OpenKnot provided participants models of secondary structure with pseudoknots in NUPACK.^47,48^ For subsequent rounds (OpenKnot Rounds 1, 2, and 3), the ThreshKnot^49^ algorithm guided by EternaFold^23^ base pair probability matrices enabled faster simulations with pseudoknots. The design interface included constant 5’ and 3’ sequences needed for experimental characterization and asked participants to design unique ‘barcode’ hairpins near the 3’ end to allow for unambiguous deconvolution of their sequences from pooled chemical mapping experiments (see next section, ‘Collation of sequence libraries’, and **Supplemental Tables S2**, **S4**, and **S9)**. Between Rounds 1 and 2, a ‘Pseudoknot detective’ round was held to collect natural molecules up to 240 in length for which literature analyses provided strong evidence for pseudoknots; and during Round 3, which was largely devoted to collecting sequences for Kaggle model evaluation, 10 pseudoknotted structures were made available as specific design targets (**Supplemental Table S2**).

An OpenKnot score was developed to provide to Eterna participants a quantitative metric for success in designing pseudoknots (https://github.com/eternagame/OpenKnotScore). The metric estimates the likelihood that the secondary structure that best fits the data has pseudoknots. Candidate secondary structures were derived from ContraFold,^50^ EternaFold,^23^ HotKnots,^51^ IPknot,^52^ iterative HFold,^53^, Knotty,^54^ NUPACK^47^ (with and without pseudoknot prediction^48^), PKNOTS,^55^ SHAPEknots,^56^ SPOT-RNA,^45^, and ViennaRNA,^57^ as well as base-pair probability matrices from CONTRAfold, EternaFold, and ViennaRNA post-processed into secondary structures with ThreshKnot^49^ and with the Hungarian^58^ algorithm after adding the probability of being unpaired as diagonal weights (implemented in ARNIE; https://github.com/DasLab/arnie). The fit of each structure to experimental SHAPE data was numerically assessed with the ‘classic’ Eterna score, ranging from 0 to 100.^21^ This score was defined as the percentage of nucleotides with reactivity < 0.5, for regions predicted to be paired, and with reactivity > 0.125 for regions predicted to be unpaired; see calc_eterna_score_classic.m available in https://github.com/eternagame/OpenKnotScore. All structures within 5 score units of the best fit structure were considered as candidate best-fit structures. The OpenKnot score, which ranges from 0 to 100, was computed as the mean of the classic Eterna score and the Eterna score computed over just nucleotides that formed pairs involved in pseudoknots, itself averaged over these best-fit structures; **Extended Data Figure 1d** shows an example OpenKnot ‘score card’ made available to Eterna participants.

### Collation of sequence libraries

For each experiment, sub-library sequences were collated and flanking sequences were added as necessary. First the sequence region of interest was prepared. All sequences had to be the same length; sequences that were longer (e.g. genomes or RFAM) were windowed with a 10 bp stride and sequences that were shorter were padded to the correct length. For M^2^-seq or other mutational scan libraries, the mutants were all added. Barcodes, unique for each sequence, were designed and added on the 3’ end of the sequence region of interest (**Supplemental Table S3** gives hairpin lengths for each library; **Supplemental Table S9** provides an example sequence). For some contributed sub-libraries the sequences were already padded and barcoded by the contributors (e.g., Eterna participants). For those that were not barcoded, barcodes and padding were designed according to the following principles. Each barcode was at least a minimum edit distance of 2 away from any other barcode in the library. To reduce misfolding of the regions of interest with the barcode regions, all barcodes were designed to be stem-loop hairpins with a 5’-UUCG-3’ tetraloop. Barcodes and pads were designed to reduce interactions with the sequence of interest, using EternaFold base-pair probabilities as predictors of these interactions. Depending on the length of the pad needed, they were designed to be unpaired, single stem-loops, or multiple stem-loops. Finally, all sequences were prepended with a constant sequence predicted to form a GAGUA-capped hairpin (5’-GGGAACGACUCGAGUAGAGUCGAAAA-3’) and appended with a constant sequence (5’-AAAAGAAACAACAACAACAAC-3’). These constant sequences were used as priming regions in chemical mapping experiments (see **Supplemental Table S9**). The code to compile these libraries can be found on GitHub (https://github.com/DasLab/big_library_design).

### Chemical mapping experiments

Oligonucleotide libraries (synthesized by Twist, Agilent, or CustomArray/Genscript; see **Supplemental Table S3**) were amplified using emulsion PCR (per reaction oil phase: 12 μL of ABIL EM90 (Evonik), 0.15 μL of Triton X-100 (Sigma Aldrich #T8787), 287.85 μL of mineral oil (Sigma Aldrich #M5904); per reaction aqueous phase: 26.625 μL of DNase/RNase-free water, 3 μL of 100 μM “Eterna” forward primer, 3 μL of 100 μM “Tail2” reverse primer (**Supplemental Table S9**), 3 μL of oligo pool template with concentration of 1 ng/μL, 1.875 μL of bovine serum albumin (20 mg/mL, Thermo Fisher #B14), and 37.5 μL of 2X Phire Hot Start II PCR Master Mix (Thermo Fisher #F125L). The oil phase mixture was chilled on ice for 30 minutes and transferred to a cold glass vial. A magnetic stir bar (Sigma-Aldrich, cat. no. Z329061) was placed into the glass vial and the stir rate set to 700 rpm; 10 μL drops of the aqueous phase mixture were added with a pipetter into the oil phase 5 times, with 10 seconds waiting period between each addition. The emulsion mixture was stirred for 10 minutes and transferred into PCR tubes for thermocycling. The thermocycler setting was 98 °C for 30 seconds; then 98 °C for 10 seconds, 55 °C for 10 seconds, and 72 °C for 30 seconds, repeated for a total of 42 cycles; 72 °C for 5 minutes, and 4 °C hold. The emulsion PCR product was purified using QIAquick PCR Purification Kit (QIAGEN, #28104).^59^

The RNA library was synthesized by *in vitro* transcription (TranscriptAid T7 High Yield Transcription Kit, Thermo Scientific #K0441) using the emulsion PCR product as the template. Each *in vitro* transcription reaction mixture contained 8 μL 25 mM nTPs, 4 μL of 5X TranscriptAid buffer, 2 μL of TranscriptAid Enzyme Mix, and 6 μL of 50 - 120 ng/μL emulsion PCR DNA. The reaction was incubated at 37 °C for 3 hours, and another 15 minutes after 2 μL of DNase I was added. The RNA was purified using RNA Clean & Concentrator-5 columns (Zymo, cat. no. R1013).

To chemically modify the RNA, the RNA was first denatured at 90 °C for 3 minutes after mixing 3 μL of 1 M pH 8.3 Na-bicine, 3 μL of 1 M KOAc, 4.65 μL of water, and 2.35 μL of 34 μM RNA. (For one experiment involving the SL5-M2seq samples prepared for 2A3 modification and corresponding no modification control, the RNA was denatured in a different buffer, mixing 2 μL of 500 mM Na-HEPES, pH 8.0, 8.65 μL of water, and 2.35 μL of 34 μM RNA.) Then 2 μL of 100 mM MgCl_2_ was added to the mixture to refold the RNA at 50 °C for 30 minutes. One copy of the RNA library was modified by 3% dimethyl sulfate (DMS), and the other copy was modified by 100 mM 2A3 ((2-Aminopyridin-3-yl)(1*H*-imidazol-1-yl)methanone, TOCRIS #7376). Each DMS modification was performed in a fume hood by adding 0.6 μL of DMS and 4.4 μL of water into the RNA, incubating at room temperature for 10 minutes, and quenching the reaction with 20 μL of 2-mercaptoethanol and 32 μL of water. For 2A3 modification, 5 μL of 400 mM 2A3 (dissolved in DMSO) was added to each reaction and incubated at room temperature for 15 minutes, and the reaction was quenched by adding 20 μL of 1 M DTT. No modification control samples were prepared by adding 5 μL of water instead of DMS, or 5 μL of DMSO instead of 2A3. The samples were purified using RNA Clean & Concentrator-5 kit.

The DMS modified RNA and corresponding no modification control sample were reverse transcribed by MarathonRT Reverse Transcriptase (Kerafast #EYU007); the 2A3 modified RNA and corresponding no modification control were reverse transcribed by SuperScript II Reverse Transcriptase (Thermo Fisher #18064022), using primers listed in **Supplemental Table S9**. Each MarathonRT reverse transcription reaction contained 10 μL of 2X Marathon buffer (100 μL stock made by mixing 40 μL of Glycerol with 10 μL of 1 M Tris-HCl, pH 8.3, 20 μL of 2 M KCl, 10 μL of 100 mM DTT, 2 μL of 100 mM MnCl_2_, and 18 μL of water), 2 μL of 10 mM dnTPs, 4.5 μL of DMS modified or no modification RNA (total mass around 1 μg, with samples diluted if necessary), 1.5 μL of 0.285 μM RTB primer, and 2 μL of MarathonRT Reverse Transcriptase. Each SuperScript II reverse transcription reaction was prepared with 4 μL of 5X SuperScript II First Strand Buffer, 2 μL of 10 mM dnTPs, 1 μL of 100 mM DTT, 1.2 μL of 100 mM MnCl_2_, 3 μL of water, 4.5 μL of 2A3 modified or no modification RNA (total mass around 1 μg, with samples diluted if necessary), 1.5 μL of 0.285 μM RTB primer, and 2.8 μL of SuperScript II Reverse Transcriptase. All reverse transcription reactions were incubated at 42 °C for 3 hours, and 6.5 μL of 100 mM EDTA and 20 μL of 0.4 M NaOH was added to each reaction and followed with an incubation at 90 °C for 3 minutes. 20 μL of acid quench mixture (7 mL stock made by mixing 2 mL of 5 M NaCl with 2 mL of 2 M HCL and 3 mL 3 M NaOAc) was added to neutralize the NaOH. The cDNA was purified using Oligo Clean and Concentrator columns (Zymo, cat. no. D4060).

Denatured polyacrylamide gel electrophoresis (10% gel in 7 M urea and 1X TBE) was used to size-select the cDNA, and the size-selected cDNA was purified by ZR small-RNA PAGE Recovery Kit (Zymo, cat. no. R1070). The purified cDNA was then amplified by PCR in a reaction prepared with 1 μL of 100 μM “cDNAamp” forward primer, 1 μL of 100 μM “cDNAamp” reverse primer (primers listed in **Supplemental Table S9**), 8 μL of water, 3 μL of cDNA template, and 12.5 μL of 2X Phire Hot Start II PCR Master Mix (Thermo Fisher #F125L). The following thermocycler protocol was used: 98 °C for 30 seconds; then 98 °C for 10 seconds, 65 °C for 10 seconds, and 72 °C for 30 seconds, repeated for 14 additional cycles; 72 °C for 5 minutes; and 4 °C hold. The amplified DNA was quantified and pooled along with 20% PhiX for Illumina next-generation sequencing either on NovaSeq X+ or (for longer sequences) MiSeq instruments (**Supplemental Table S3**). One library of test sequences (Positives390), which could not be probed until after the competition and gave very poor signal-to-noise (**Supplemental Table S4**), was not included in analyses presented here except studies of sequence dropout.

Sequencing data FASTQ files were analyzed using an Ultraplex-Bowtie2-RNAFramework pipeline,^60–62^ with mutations and deletions, but not insertions, counted towards chemical mapping signals (https://github.com/DasLab/ubr). The positions of deletions in same-nucleotide stretches are ambiguous; these signals were distributed across same-nucleotide stretches in direct proportion to the observed mutation signals (which are not ambiguous). The profiles were background subtracted based on the no modification control experiments and normalized so that the 90th percentile reactivity across all probed sequences was set to 1.0. Data in regions at very 5’ and 3’ ends could not be recovered as mutations were replaced by constant primer sequences. Errors were estimated based on counting statistics and propagated to final normalized profiles using standard formulae; to ensure that error estimates remained above zero, a pseudocount of 1 was added for error estimates. Background-subtracted data at 3’ barcode hairpins had high estimated errors in some libraries and were not included in any of the final profiles. Signal-to-noise ratio estimates were made based on the mean value of the reactivities at all positions with non-zero reactivity, divided by the mean value of the estimated errors at the same positions; the first and last positions with non-zero reactivities were ignored if at least 4 such positions were available.

### Ribonanza Kaggle competition

The Kaggle competition (https://www.kaggle.com/competitions/stanford-ribonanza-rna-folding/) involved preparation of several data sets and files. Training data (see **Supplemental Table S4**) were made available as profiles, giving the chemical probe type (2A3 or DMS), the reactivities at each probed position (left blank for unprobed positions), experimental errors estimated for each position, profile-level signal-to-noise estimates, total number of Illumina reads for each profile, and a simple Boolean flag to annotate higher quality profiles (SN_filter, set to 1 if the profile had signal-to-noise > 1.00 and reads > 100 and 0 otherwise). A text file of the test sequences for which 2A3 and DMS reactivity predictions were to be submitted was also provided, along with a sample submission file. Additional files provided to participants were sequence libraries grouping all 2,150,401 sequences, with titles, into the actual collections that were synthesized together; secondary structure predictions from various packages that had been compiled for OpenKnot scoring (see above); files of Eterna ID, Eterna author, title, description, and sequence for Eterna OpenKnot sequences; files listing pairs of positions predicted to have non-zero Watson-Crick-Franklin base pair probabilities by the LinearPartition package^63^ with EternaFold parameters^23^; 3D coordinates predicted by RhoFold^64^ for a subset of the training sequences with SN_filter = 1; and 67,000 previously available chemical mapping profiles from the RNA Mapping DataBase,^65^ compiled into a single file with format matching the main training data file (**Supplemental Table S10**).

### Training Kaggle models, RNAdegformer, and RibonanzaNet

Top-ranking models prepared by Kaggle competitors were prepared with a diverse set of architectures and training protocols; brief descriptions, including estimates of computational costs and links to code and more detailed summaries, are provided in **Supplemental Table S5**. Before the Kaggle competition, baseline studies were performed with RNAdegformer. RNAdegformer uses a series of 1D convolution and deconvolution in conjunction with Transformer encoder modules on the sequence representation of RNA and processes 2D features of BPP matrices stacked with inverse distance matrix using deep residual 2D convolution layers, which are used as attention biases in each Transformer encoder layer (**Fig. 2a**). Studies of data scaling and sublibrary ablation used 167,671 sequences that achieved signal-to-noise greater than 1.0 in both DMS and 2A3 experiments, with 139,725 sequences (83%) split out for training while reserving the rest for validation. When using BPP as a feature, 256 hidden states were used for RNAdegformer and when only using sequence as input, 512 hidden states were used. When using BPP as a feature, RNAdegformer was trained for 30 epochs, using the Ranger optimizer^66^ on a flat and anneal learning rate schedule, with learning rate starting at 0.001. When only using sequence as input, RNAdegformer was trained for 120 epochs in data scaling experiments and for 70 epochs in dataset dropout experiments. The final RNAdegformer model that served as a baseline for the Kaggle competition was also trained with sequences that had at least one profile of 2A3/DMS with high signal to noise ratio (the other profile was masked during gradient computations to update the network), resulting in a total of 214,831 training sequences, and trained for 30 epochs. This model took 5.4 hrs to train on two NVIDIA RTX 3090 GPUs.

After the Kaggle competition, a new architecture called RibonanzaNet was developed, which does not use BPP features but instead creates a fully learned pairwise representation (Fig. 4a). RibonanzaNet bears some similarities to top-ranking Kaggle models and RNAdegformer, because it combines 1D convolutions with Transformer encoder modules. However, the pairwise representation is updated globally, unlike BPP features used in RNAdegformer, which can be seen as a pre-computed pairwise representation. Following an embedding layer that transforms RNA bases into sequence representation, RibonanzaNet spawns a pairwise representation by computing pairwise outer products from a downsampled sequence representation. Then relative positional encodings up to 8 bases apart are added to the pairwise representation. Next, RibonanzaNet processes the sequence and pairwise representation through several layers via 1D convolution, self-attention, and triangular multiplicative updates. The combination of 1D convolution and self-attention allows the model to learn interactions between RNA bases or short segments of bases (*k*-mers) at any sequence distance, while leveraging information in the pairwise representation. Further, the outer product mean operation updates the pairwise representation using projected outer products, and triangular multiplicative update modules operate on the pairwise representation to update each edge with two other edges starting from/ending at the two nodes of the edge being updated. It is important to note that while the RNAdegformer and other Kaggle models that use BPP features to bias self-attention have information flowing only from BPP representation to sequence representation, in RibonanzaNet, information flows not only from the pairwise representation to sequence representation but also from sequence representation back to pairwise representation.

The training procedure for RibonanzaNet went through multiple stages (**Extended Data Fig. 7a**). An initial model was trained using sequences that have either or both 2A3/DMS profiles with signal to noise ratio above 1.0 (214,831 training sequences) over 30 epochs, which took 11 hours on 10xL40S GPUs. A second model ensembled predictions for the remaining (noisy) training data from the top 3 Kaggle models as pseudo-labels (563,796 sequences), and pre-trained RibonanzaNet for 30 epochs with these pseudo-labels and a flat learning rate of 0.001, followed by 10 epochs of training solely on true labels of training sequences that have one profile of 2A3/DMS with high signal-to-noise ratio, following a cosine learning rate schedule that annealed learning rate to 0. This entire protocol took 30 hours on a server with ten NVIDIA L40S GPUs. A final model ensembled predictions of top 3 Kaggle models for noisy labels from training data as well as for all test sequences to prepare a new pseudo label dataset of 1,907,619 sequences. Repeating RibonanzaNet training on this larger set of pseudo-labels and then annealing on true training labels for high signal-to-noise data took 140 hours on 10xL40s GPUs. Sequence flip augmentation was used throughout training; a model trained without this augmentation in the final ‘annealing’ gave worse MAE test accuracy. None of these RibonanzaNet versions made use of experimental data labels for the test sequences.

### Predicting sequence dropout in chemical mapping experiments

The task of fine-tuning RibonanzaNet to predict sequence dropout required converting the one-dimensional encoding outputted by the network to a pair of positive numbers per sequence, the number of 2A3 and DMS read counts obtained at the end of the experiment. This was accomplished by passing the model embeddings through a dense layer, taking the mean over the entire sequence, and passing the result through an exponential. The model was fine-tuned by training on the experimental number of sequencer read counts from the DasLabBigLib-1M dataset and tested on sequencer read counts for two other sequence libraries with different lengths, DasLabBigLib2-1M and Positives240 **(Supplemental Table S3)**. Since the training and validation data spanned multiple orders of magnitude and empirically followed a log-normal distribution, the MSE of log-reads was taken to be the loss function used for training and validation. To bring the predictions and data to the same scale, the read counts for both were scaled so that the average number of reads per sequence was a fixed value (1,000, the target number of average reads in our experiments).

During the fine-tuning process, overfitting was prevented by scheduling the fine-tuning process, progressively unfreezing the layers of the model one epoch at a time, with the first epoch reserved for training the dense layer to avoid distorting the pre-trained features. The Adam optimizer, with a learning rate of 0.001 scheduled over the first ten epochs, was employed. Overfitting was further reduced by employing early stopping and selecting the model with the lowest validation loss. Since the prediction of library dropout has been given little attention in the literature before, a set of alternative models was trained for comparison. The same fine-tuning process for RibonanzaNet-Drop was carried over to the RNA-FM foundation model,^40^ and in addition two LSTM models were trained from scratch – one which had access only to the sequence, and another which was fed both the sequence and predicted secondary structure from ViennaRNA^57^ in dot-bracket form. The training specifics were all kept consistent with those used for RibonanzaNet fine-tuning. As additional simple baselines corresponding to common heuristics used to avoid sequence synthesis issues, Bayes estimators were fitted using the length of the longest repeat in the sequence and the length of the longest hairpin predicted for that sequence in ViennaRNA.

### Fine-tuning RibonanzaNet to predict RNA hydrolytic degradation

RibonanzaNet was fine-tuned to predict the nucleotide-resolution properties characterizing RNA hydrolytic degradation in the OpenVaccine challenge: reactivity to SHAPE modified 1M7; hydrolysis in the presence of 10 mM MgCl_2_ and pH 10.0 buffer at 25 °C; and hydrolysis in the presence of 10 mM Cl_2_ and pH 8.0 buffer at 50 °C. Because this task is similar to predicting chemical reactivity of 2A3/DMS experiments, the last linear layer of RibonanzaNet was simply replaced with a new one that outputs multiple predictions per position. Since the OpenVaccine dataset contains many sequences with low signal to noise, sequences whose signal-to-noise values were less than 1.0 were excluded from training. RibonanzaNet-SS was then fine-tuned for 20 epochs with the Ranger optimizer^66^ using RMSE loss with a flat and anneal learning rate schedule.

### Fine-tuning RibonanzaNet to predict secondary structure

RibonanzaNet was fine-tuned to predict secondary structure using secondary structures derived from known 3D RNA structures in the PDB.^17,66^ Train/test splits for this PDB dataset were derived from reference^11^ and further filtered by removing duplicate sequences, duplicate structures, structures that contained no base pairs, and structures with non-canonical nucleotides. Following this filtering, the PDB dataset was left with 631 training sequences and 66 test sequences. All 12 RNA sequences from CASP15 were included as a second test set. Data are provided in **Supplemental Table S8**.

As secondary structure prediction fundamentally involves predicting interactions between nucleotides, a linear layer was added on top of the final RibonanzaNet pairwise representation to make predictions. The resulting RibonanzaNet-SS model was then fine-tuned with binary cross entropy loss on the ground truth 2D connectivity matrix *M* representing secondary structure, where pairing between nucleotides *i* and *j* is represented as *M_i,j_* = 1 and all other entries within the matrix are 0. Positive labels were up-weighted by 2, due to the sparsity of positive labels. Training loss contributions from sequences were weighted by the square of their lengths to account for the larger amount of information in long secondary structures. Fine-tuning for secondary structure was performed based on two versions of RibonanzaNet: one network that had been pre-trained using chemical mapping data and one with the same architecture without pre-training on chemical mapping data. For both networks, an initial learning rate of 0.0001 was used along with a cosine annealing learning rate schedule. The pre-trained network was trained for a total of 5 epochs and the network without pre-training for a total of 20 epochs, taking the model parameters from the epoch with the lowest validation loss for further evaluation. Batch size for all fine-tuning was 1. Following fine-tuning, the Hungarian algorithm as implemented in ARNIE was used to create a base-pairing matrix where each nucleotide only has one partner. Entries with values lower than 0.5 were ignored, based on studies varying this threshold between 0 and 1 and using a random subset of the training set for validation. Base pairs *i-j* with |*i – j|* < 4 (i.e. hairpins with loop lengths of 0, 1, or 2) were filtered out. ‘Lone’ base pairs (singlet stems), which have been filtered out in some other studies, were retained to allow evaluation of tertiary structure signatures like Levitt base pairs.^67^ The final matrix of base pairs was then compared against labels in the test set to score predictions for precision, recall, and *F*_1_ (harmonic mean of precision and recall). To match prior studies, test sequences for the PDB test set (but not the CASP15 dataset) were clustered according to the sequence similarity using the same clusters as in Boyd et al.^9^ (*test_cluster_idx* in Supplemental Table S8) and we report statistics using the mean *F*_1_ score from each cluster. RibonanzaNet-SS modeling accuracy did not depend on the length of test sequences, structural homology or sequence homology to training data (**Extended Data Fig. 13**).

To more specifically evaluate pseudoknots which involve non-nested or ‘crossed’ pairs (base pairs *i*-*j* and *m*-*n* with *i* < *m* < *j* < *n*), we also computed a metric *F*_1.crossed-pair_, taken as the harmonic mean of precision and recall after filtering both the reference and modeled base pair list for just crossed pairs. Targets for which both the reference and modeled secondary structure were pseudoknot-free (no crossed pairs) were not included in computing mean or median summary statistics (blank in **Supplemental Tables S8**). However, on targets for which the reference had no crossed pairs but the predicted structure did include crossed pairs, a value of *F*_1.crossed-pair_ = 0 was assigned, i.e., overprediction of pseudoknots was penalized.

The predicted pair scores *M_i,j_* outputted by the fine-tuned RibonanzaNet typically reside between 0 and 1; as an estimate of modeling confidence, the mean of the pair scores over the final secondary structure <*M_i,j_*>_ss_ was computed. The secondary structure prediction accuracy and model confidence appeared to be linearly correlated, and confident predictions tend to be more accurate. A univariate linear regression model was fitted to these data to estimate *F*_1_ score of predicted secondary structure (**Extended Data Figure 9a**; fitted relation *eF*_1_ = 2.25 <*M_i,j_*>_ss_ – 1.29). A regression model was also fitted that estimates *F*_1_ score of predicted pseudoknots using average predicted probabilities of crossed base pairings (**Extended Data Figure 9b**; fitted relation: *eF*1.crossed-pair = 2 <*Mi,j*>crossed-pair – 1.19).

### Three-dimensional RNA structure modeling with trRosettaRNA

trRosettaRNA source code and model weights were downloaded from https://yanglab.qd.sdu.edu.cn/trRosettaRNA/download/ or requested from trRosettaRNA authors. trRosettaRNA utilizes a base-pairing probability matrix in combination with an MSA and sequence features to generate 3D predictions. MSAs were derived from the trRosettaRNA server (https://yanglab.qd.sdu.edu.cn/trRosettaRNA/). 3D predictions were generated by trRosettaRNA utilizing either the default BPP matrix predicted by SPOT-RNA^45^ or the pair score predicted by RibonanzaNet-SS. For CASP15 test structures, model weights downloaded on October 18th, 2022 were used to generate predictions based on training data that approximated structures available for CASP15^3^. The PDB training and test data developed for RibonanzaNet-SS contains structures that were published before April 30, 2020; trRosettaRNA model weights from April 2019 were therefore used to reduce train/test leakage when generating predictions for the PDB test set. The trRosettaRNA folding calculations were carried out using default parameters to generate 5 models. The model with the lowest Rosetta energy score was then selected for further evaluation. RMSD and lDDT values were computed with rna-tools^68^ (https://github.com/mmagnus/rna-tools) and OpenStructure^69^ (https://openstructure.org), respectively. For statistical analysis, we first calculate the lowest RMSD or highest lDDT value of each trRosettaRNA prediction against all available models for a test molecule. The lowest RMSD or highest lDDT value per cluster (defined by the same *test_cluster_idx* used for secondary structure evaluation) is reported to avoid overweighting similar structures in the analysis. Links to files of predicted and reference structures and code to reproduce 3D analysis are available in **Supplemental Table S10.**

## Data availability

Datasets including Ribonanza chemical mapping profiles, raw Illumina sequencing data, and RibonanzaNet models are available on RNA Mapping Database, the Sequence Read Archive, Kaggle, Github, and Google Drive at links provided in **Supplemental Table S10**. Post-competition Ribonanza+ data are freely available for non-commercial use at https://eterna.tech.

## Code availability

Code including library preparation scripts, data processing scripts, and notebooks for neural network training and inference are available on Kaggle and GitHub at links provided in **Supplemental Table S10**.

## Extended Data

**Extended Data Figure 1.**
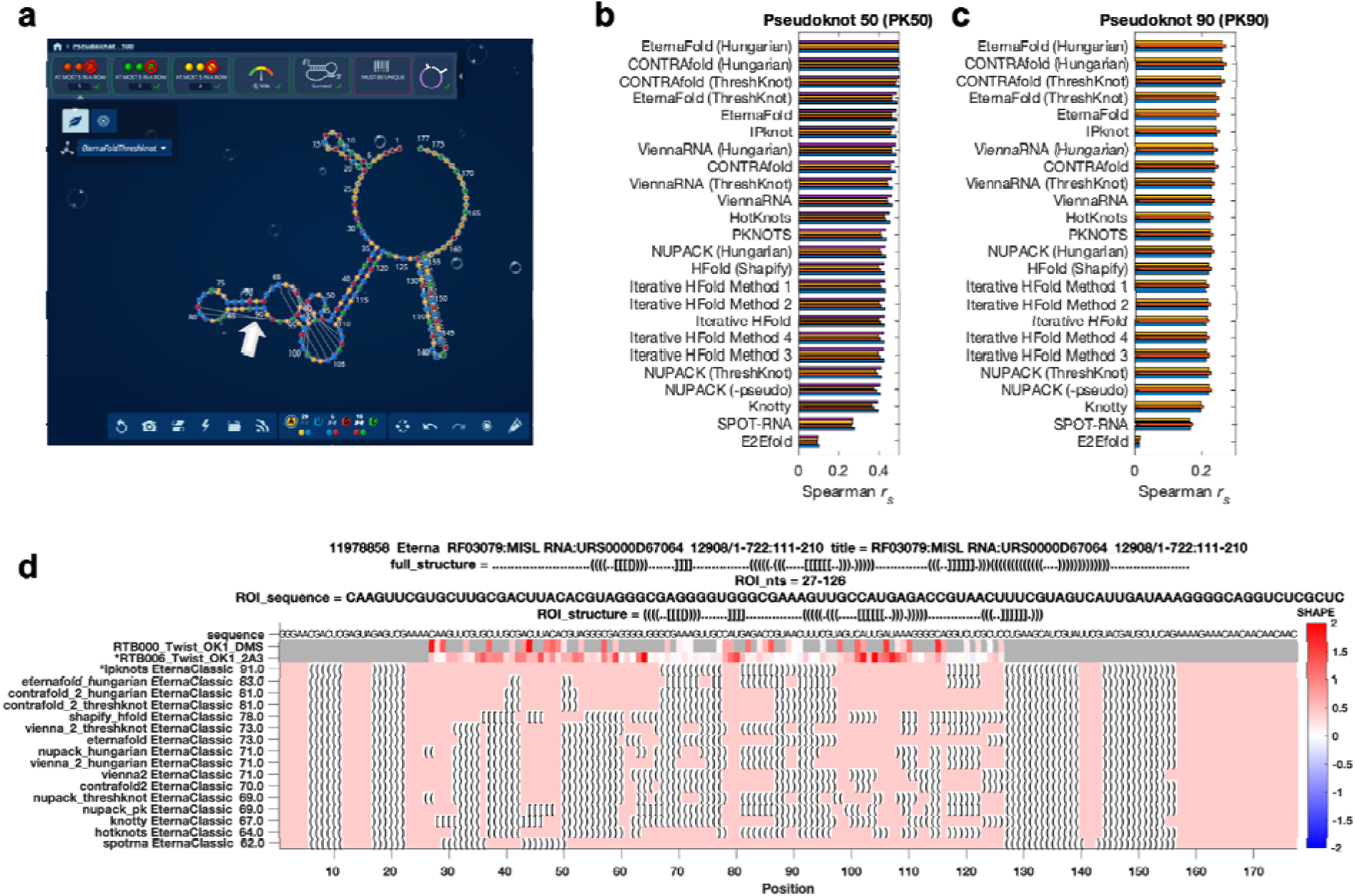
Eterna OpenKnot challenge. **(a)** Eterna interface, showing straight strings (white arrow) to mark pseudoknots modeled with EternaFold and ThreshKnot. **(b-c)** Accuracy of secondary structure modeling packages. Paired and unpaired nucleotides in secondary structure predictions were converted to 0 and 1; correlation coefficients (Spearman *r_s_*) were computed against SHAPE (2A3) data from two separate libraries with insert lengths of **(b)** 50 and **(c)** 90 nucleotides from Eterna OpenKnot pilot rounds; different colored bars show results from four and three replicates, respectively. Figure is limited to single-structure comparisons since most packages for pseudoknot prediction do not model structural ensembles. Data are derived from experiments on PK50 and PK90 listed in **Supplemental Table S10**. **(d)** Example of design card made available to Eterna players: chemical mapping data derived from DMS and 2A3 mapping experiments (top tracks) allow ranking of secondary structures (bottom tracks; ipknots = IPknot) predicted for a window of a MISL RNA from RFAM.

**Extended Data Figure 2.**
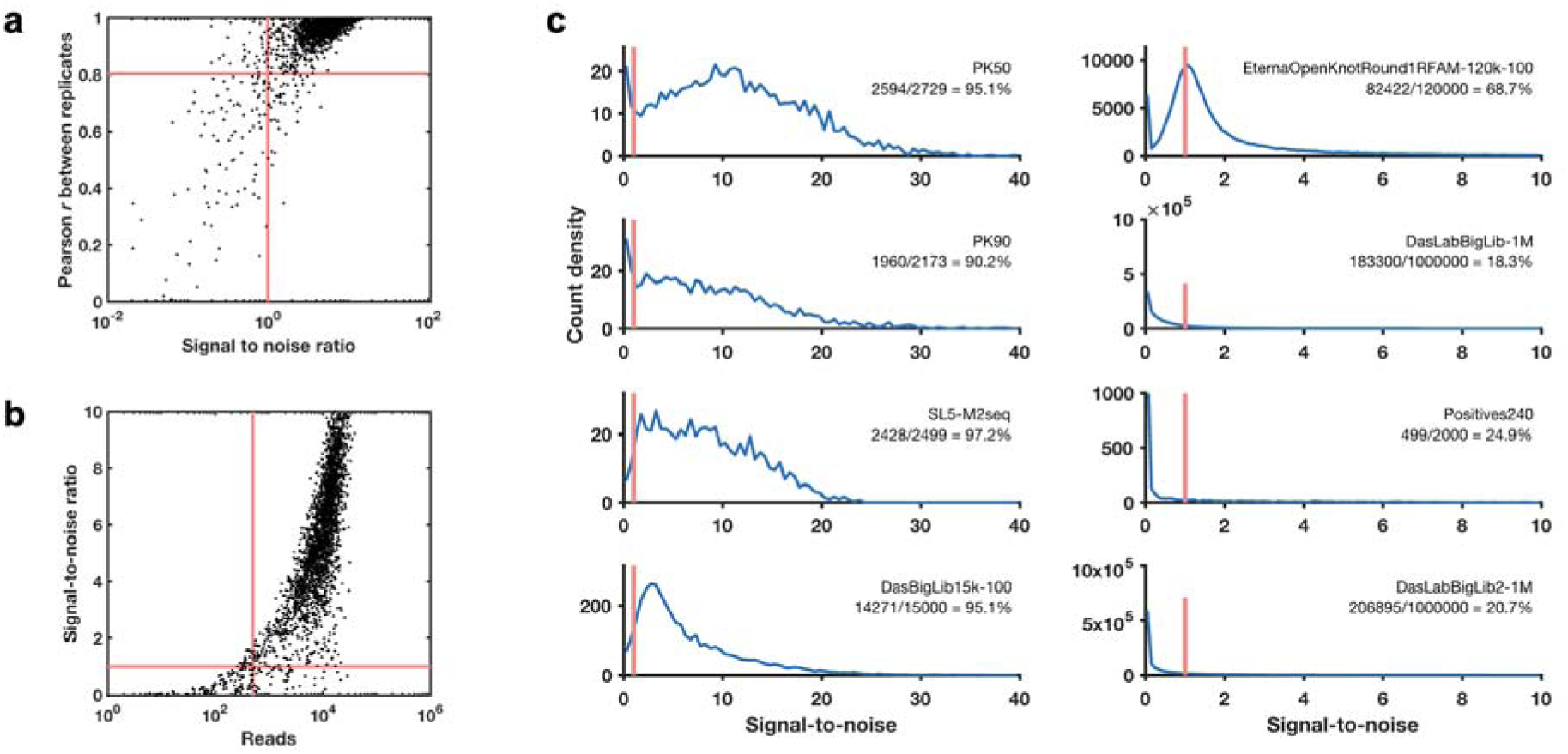
Signal-to-noise across Ribonanza data sets. **(a)** Signal-to-noise ratio values estimated based on Illumina counting statistics predicts replicability as assessed by Pearson correlation coefficient *r* between replicate datasets; a signal-to-noise ratio of 1.0 corresponds to *r* = 0.80 (red lines). **(b)** The number of reads correlates with signal-to-noise ratio, with a read number of 500 corresponding to a mean signal-to-noise ratio of 1.0 (red lines). **(c)** Experiments that seek data on larger numbers of sequences or longer sequences (‘Positives240’, with insert length of 240 compared to 50-130 nucleotides) give smaller fractions of sequences with signal to noise ratio above 1.0 (red bars). Note shift in x-axis scale in right-hand four panels compared to left-hand four panels. In all panels, results for SHAPE profiles with the 2A3 modifier are shown. In **(a)-(b)**, replica datasets were experiments for the Eterna OpenKnot Pseudoknot 50 (PK50) pilot datasets carried out with DNA prepared by two different synthesis companies (GenScript, Twist) by two different experimenters (P50LIB_2A3_000001, P50LIB_2A3_000002 in **Supplemental Table S10**).

**Extended Data Figure 3.**
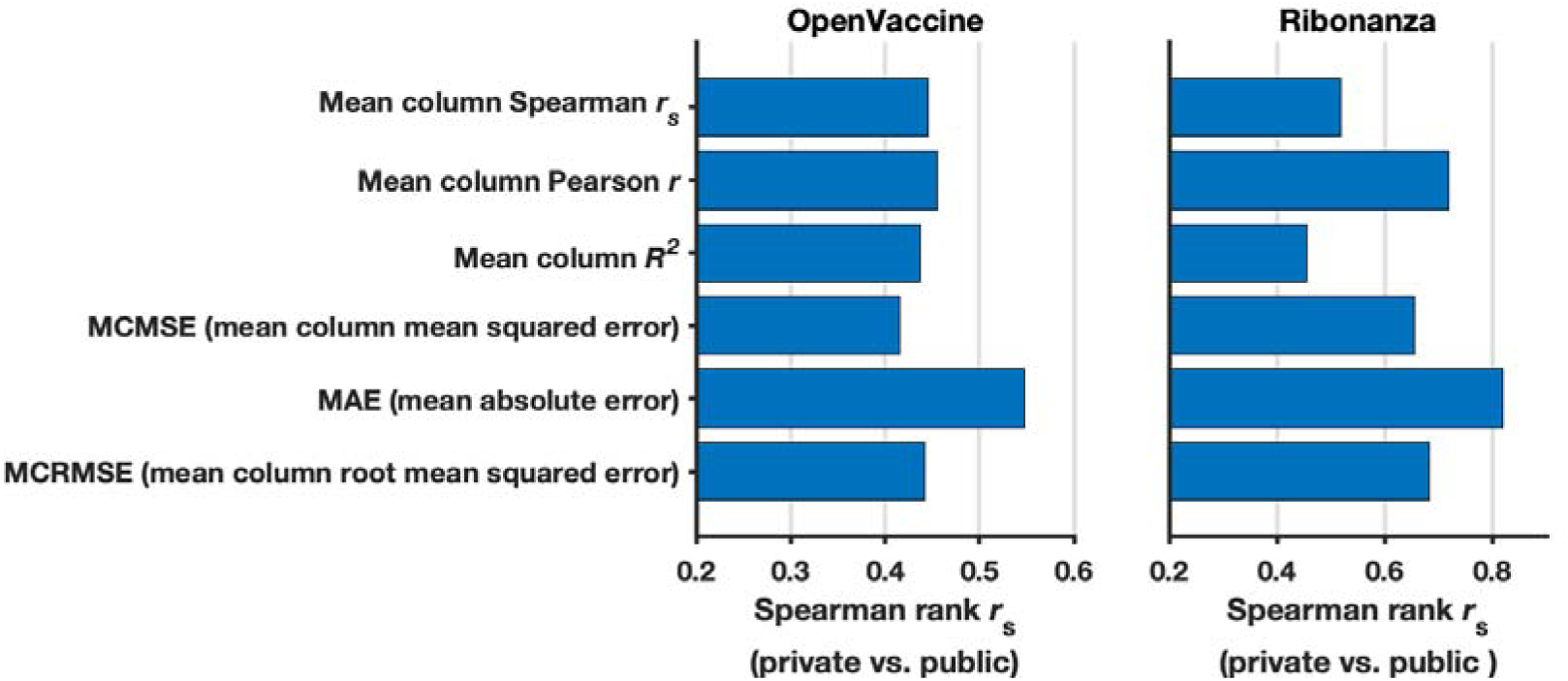
Rationale for choosing mean absolute error (MAE) as evaluation metric for Ribonanza Kaggle competition. **(a)** To determine what metric to use for scoring Kaggle submissions against experimental data, we rescored top 10 public/private submissions from the preceding OpenVaccine Kaggle competition (left) to see which metric resulted in the least shakeup between public leaderboard scores (test data for which participants could see scores but not individual data for continuous evaluation) and private leaderboard scores (test data completely unavailable to participants), as measured by Spearman *r*_s_ between public/private scores. MAE was the best in preventing shakeup (highest Spearman *r*_s_ between public/private scores). Consistent with OpenVaccine competition scoring, data were not clipped for OpenVaccine comparisons. **(b)** We rescored top 10 public/private submissions from the Ribonanza competition and confirmed that MAE had the highest Spearman *r*_s_ between public/private scores (least shakeup). Consistent with Ribonanza competition scoring, data were clipped between 0 and 1 for Ribonanza comparisons.

**Extended Data Figure 4.**
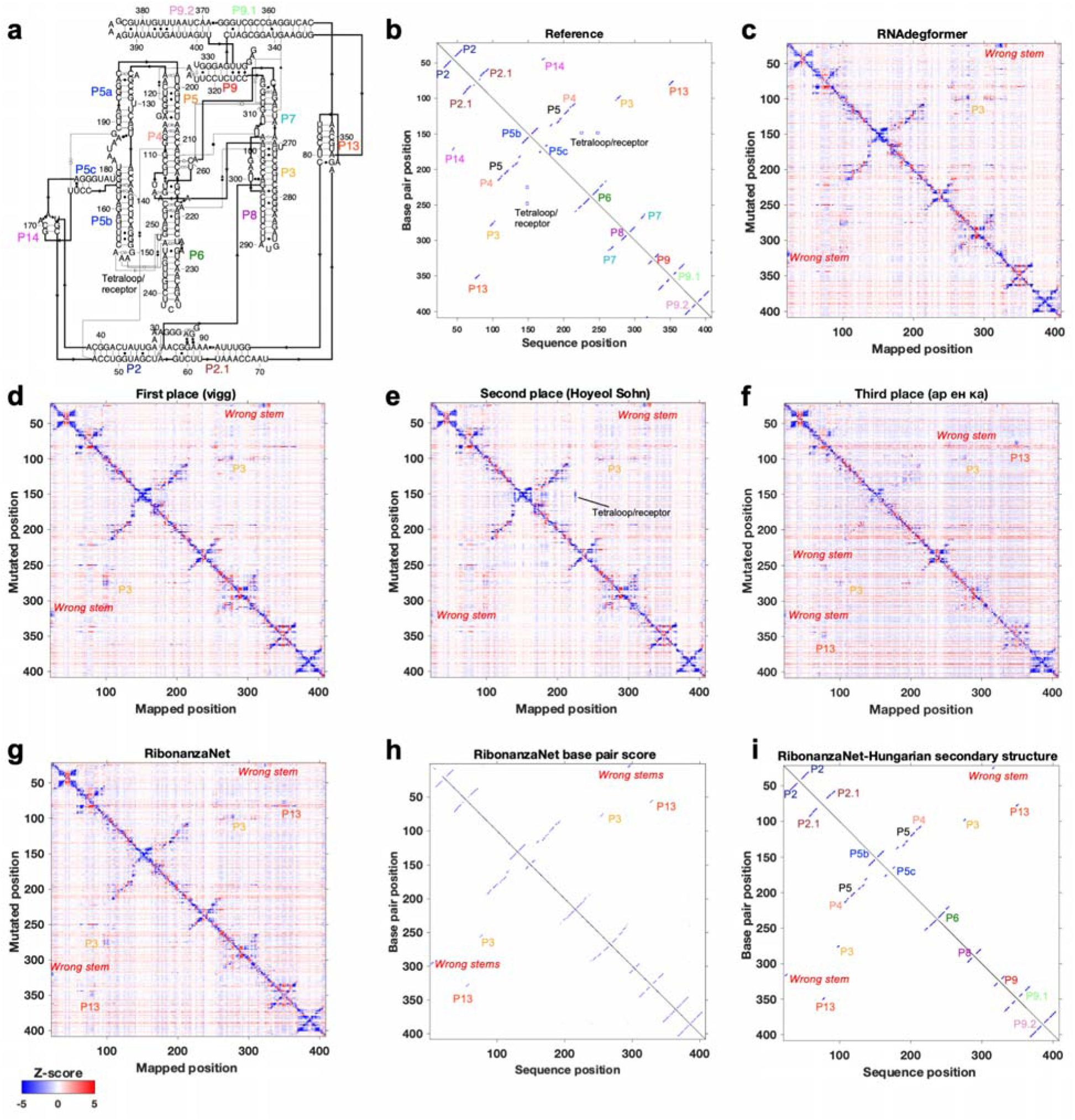
Ribonanza results on *Tetrahymena* ribozyme. **(a)** Secondary structure and **(b)** 2D map of stems and tetraloop/receptor tertiary contact inferred from cryo-EM structure (PDB: 7EZ0). M^2^ predictions from **(c)** RNAdegformer baseline, **(d-f)** Kaggle 1st, 2nd, and 3d place, and **(g)** RibonanzaNet models mark out stems in the molecule, including a P3 pseudoknot in the RNA’s catalytic core (gold), but different models predict different spurious stems (red labels), and all except Kaggle 2nd place model miss the tetraloop/receptor. **(h)** RibonanzaNet base pair score leads to **(i)** mostly accurate secondary structure prediction (*F*_1_ = 0.85) whose inaccuracy at pseudoknots is flagged by RibonanzaNet’s estimated accuracy value *eF*1,crossed pair = 0.46.

**Extended Data Figure 5.**
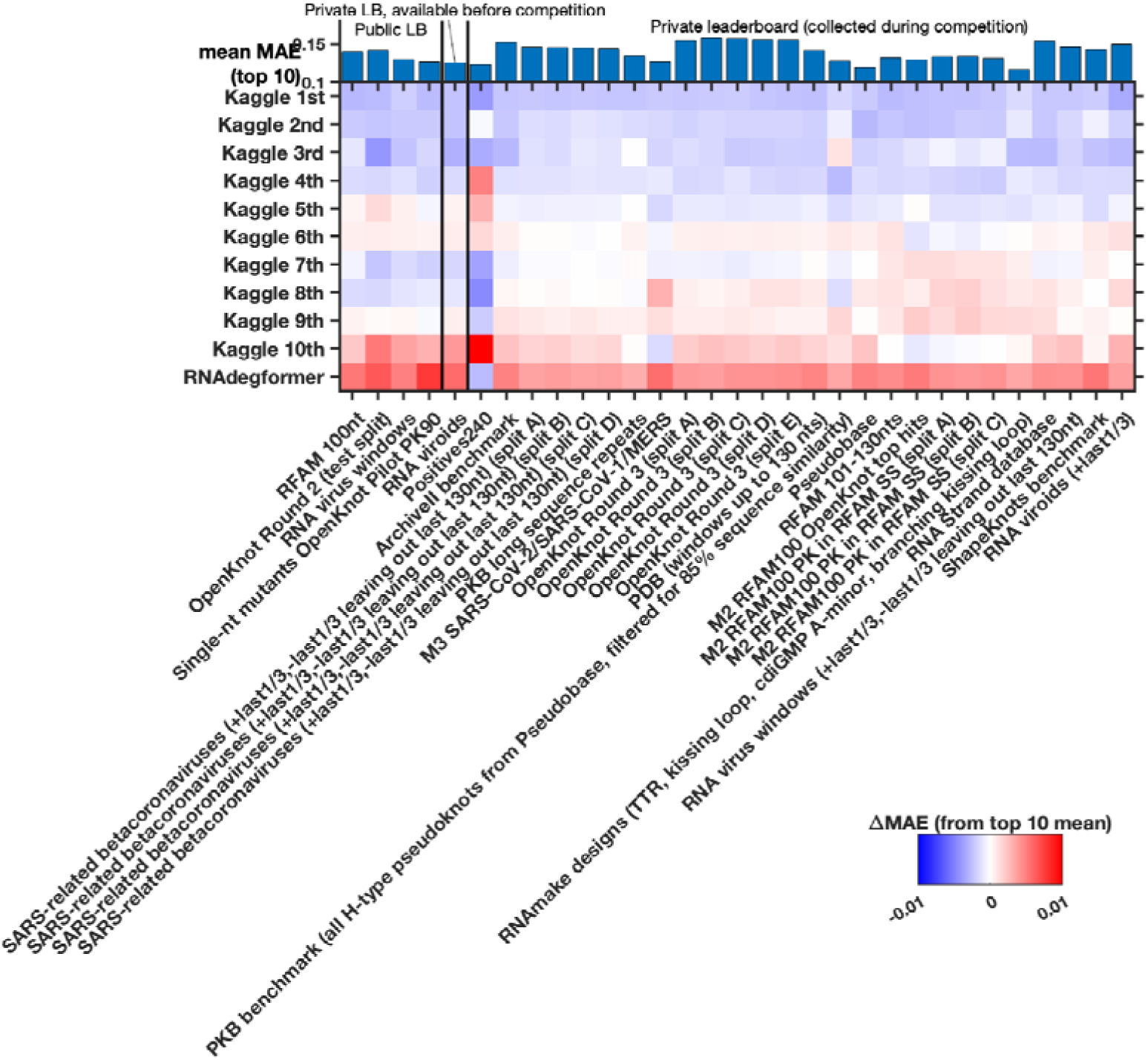
Different Kaggle models perform best for different test sub-libraries of the test set. Heatmap gives MAE accuracy to experimental data (here presented relative to mean MAE over top 10 models, shown in the top bar graph). Some of the larger sub-libraries were split (‘split A’, ‘split B’, etc.) to simplify data processing.

**Extended Data Figure 6.**
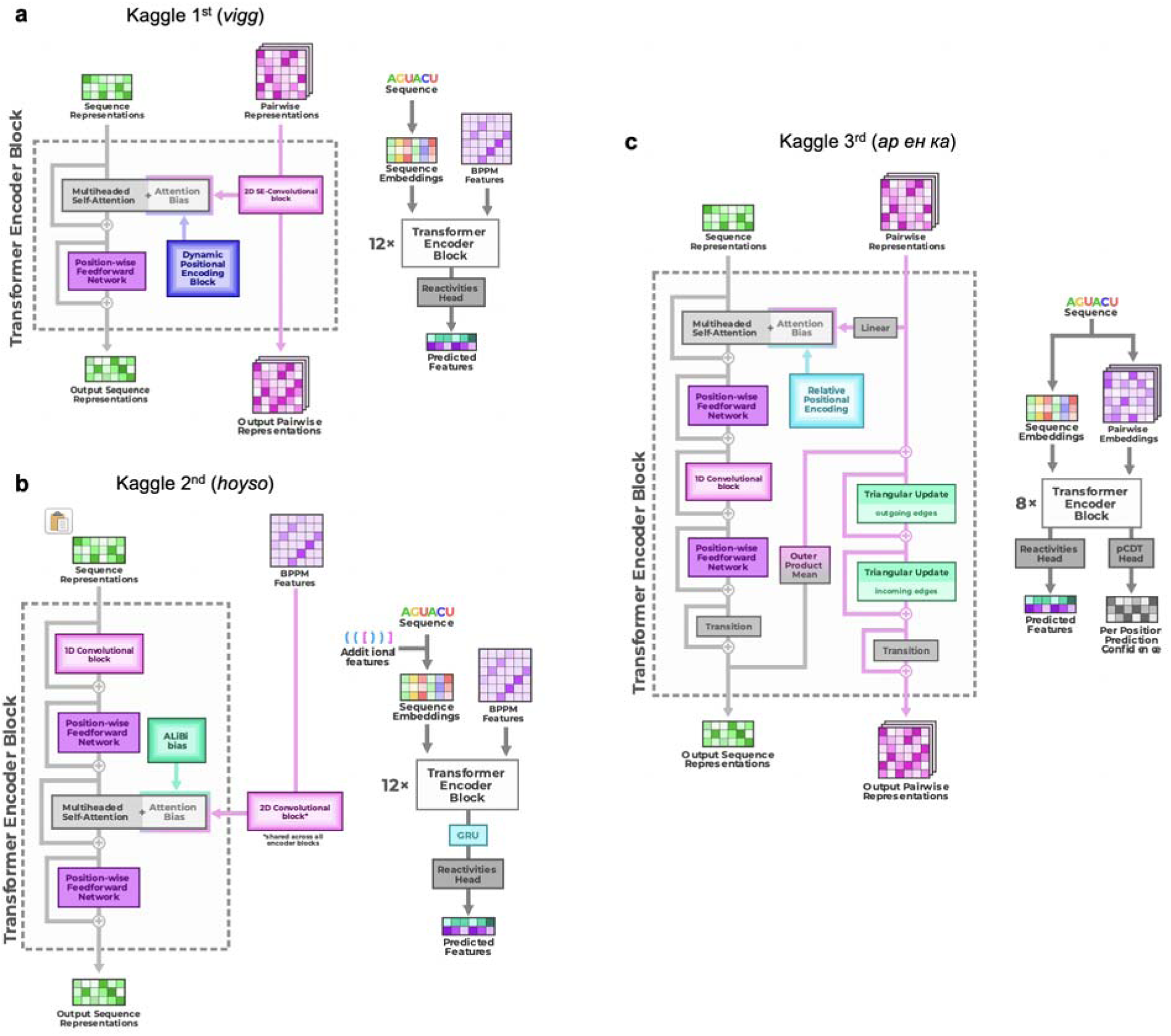
Full architecture diagrams of top 3 Ribonanza Kaggle models. **(a)** 1st place model (team *vigg*), **(b)** 2nd place model (team *hoyso*), and **(c)** one of two models used for 3rd place submission (Twin Tower model from team *ар ен ка*).

**Extended Data Figure 7.**
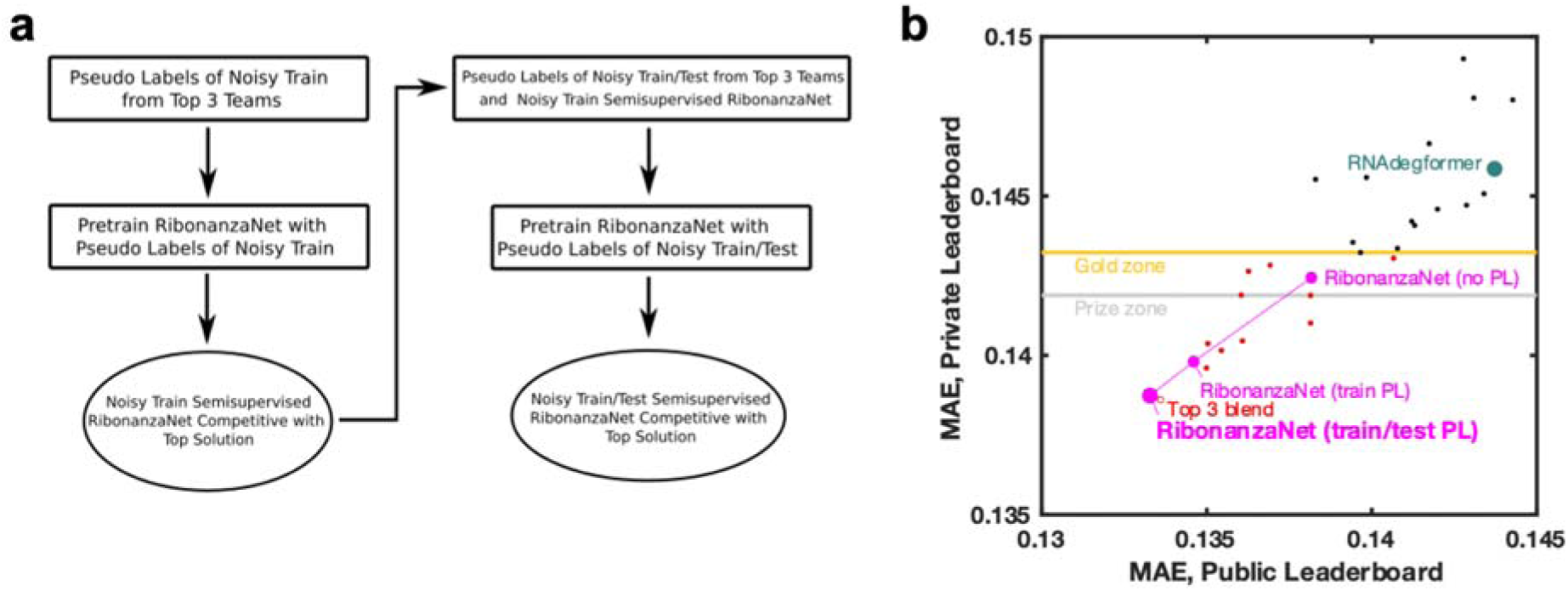
Training RibonanzaNet. **(a)** Steps taken to train RibonanzaNet, initially with pseudo labels from top 3 Kaggle submissions over the train sequences with noisy data; then training with pseudolabels expanded to include test data; and finally ‘semisupervised’ learning including actual data for train sequences. **(b)** improvements of RibonanzaNet test accuracy (MAE, mean absolute error to test data after clipping values between 0 and 1) as more pseudo-labels were included. ‘Gold zone’ and ‘prize zone’ mark 11th place and 6th place Kaggle scores which were cutoffs for Kaggle gold medals and prizes, respectively.

**Extended Data Figure 8.**
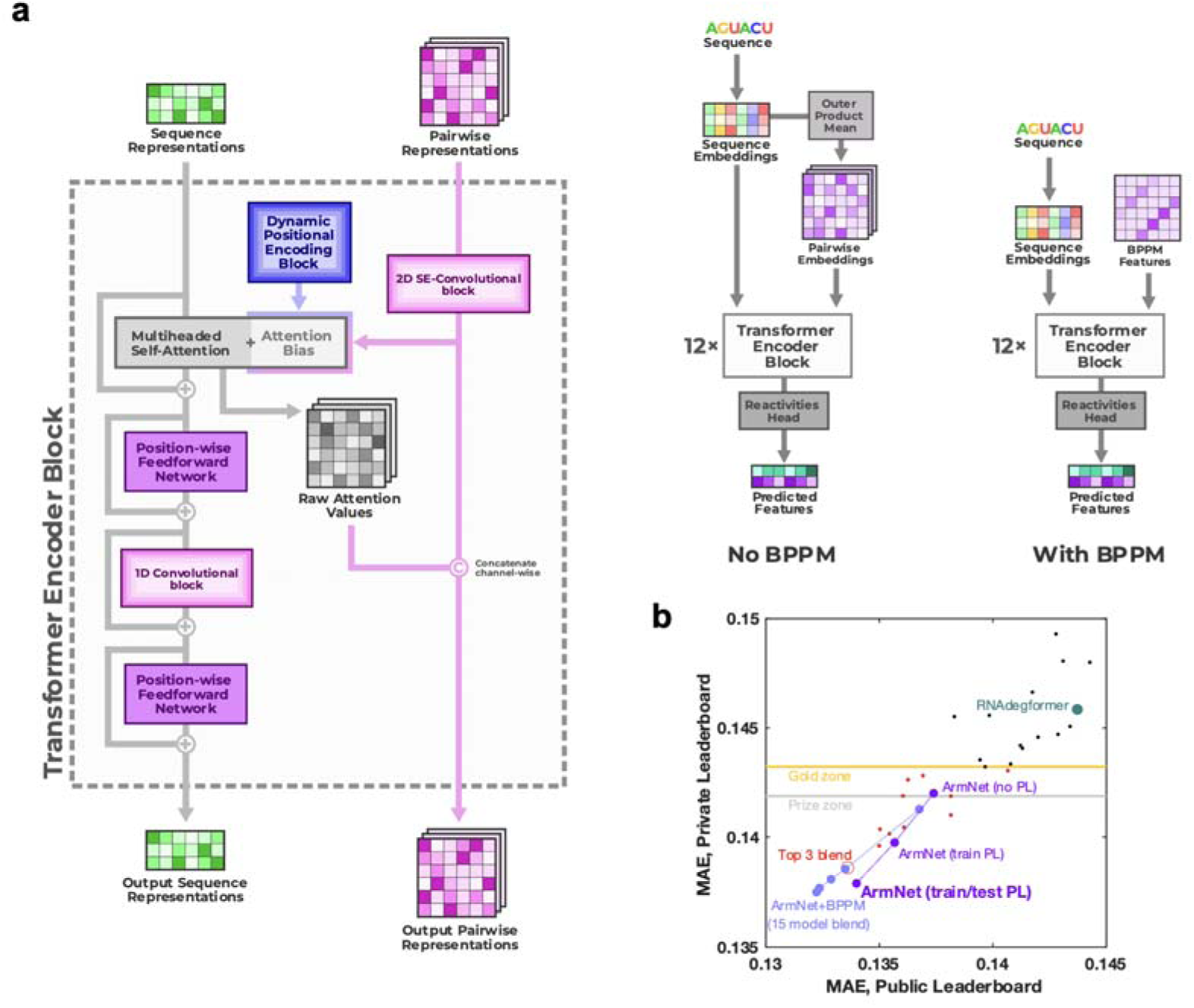
ArmNet (Artificial Reactivity Mapping using neural Networks) post-competition model from the *vigg* team. **(a)** Two modifications to the Kaggle 1st place model improved performance: (1) adding the 1D convolutional module after each attention block, as was done in the Kaggle 2nd place solution, and (2) concatenating the attention scores and BPP features and combining them using the 2D convolutional layer of the next block. The second modification supports the idea from RibonanzaNet and the 3rd place solution – to provide two-way communication between the sequence and 2D features – is important for model performance. Triangular operations tested in RibonanzaNet were not included in ArmNet. **(b)** With input of BPP matrices from EternaFold and blending an increasing number of models (1, 3, 5, 7, 15), ArmNet outperforms all previous models in both private and public leaderboard MAE (light blue symbols). Without BPP and as a single model, ArmNet achieves excellent private leaderboard MAE when trained on pseudo labels from the 15-model ArmNet-BPPM ensemble (‘PL’; purple). When the single no-BPP ArmNet model is instead trained on pseudolabels derived from the top 3 Kaggle submissions (as in RibonanzaNet; **Extended Data Figure 7**), MAE scores are slightly worse than other ArmNet models and RibonanzaNet (not shown; see **Supplemental Table S5**). MAE is mean absolute error to test data after clipping values between 0 and 1. ‘Gold zone’ and ‘prize zone’ mark 11th place and 6th place Kaggle scores which were cutoffs for Kaggle gold medals and prizes, respectively.

**Extended Data Figure 9.**
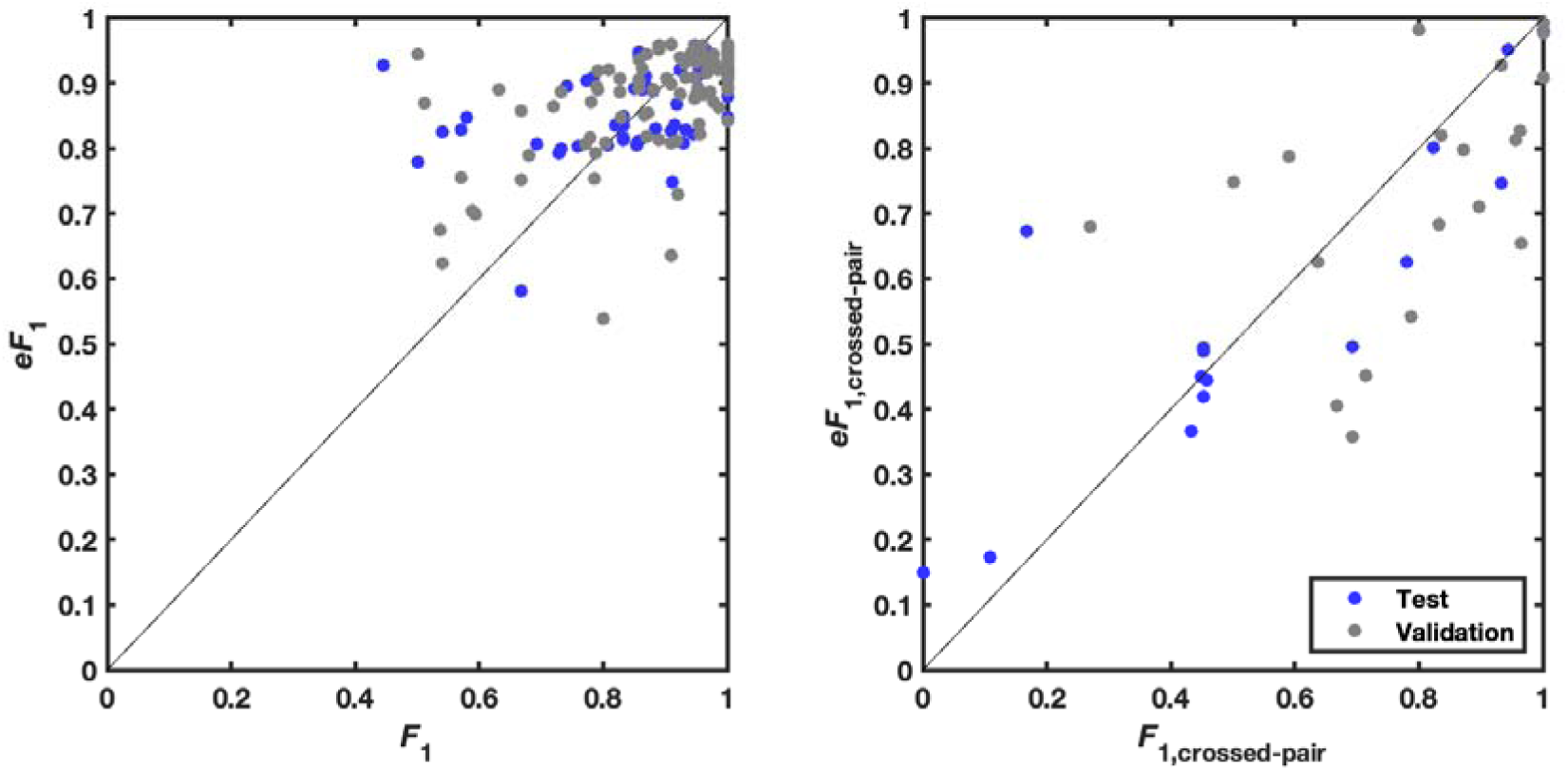
Estimation of confidence in secondary structure modeling. Expected *eF*_1_ (harmonic mean of base pair precision and recall) vs. actual *F*_1_ over (a) all base pairs and (b) just base pairs in pseudoknots (pairs *i*-*j* that ‘cross’ another pair *m*-*n*, i.e., *i* < *m* < *j* < *n* or *m* < *i* < *n* < *j*). Values for secondary structures in the test set as well as a random held out split of the train set (‘validation’), which were not used to fit the *eF*_1_ relations, are shown.

**Extended Data Figure 10.**
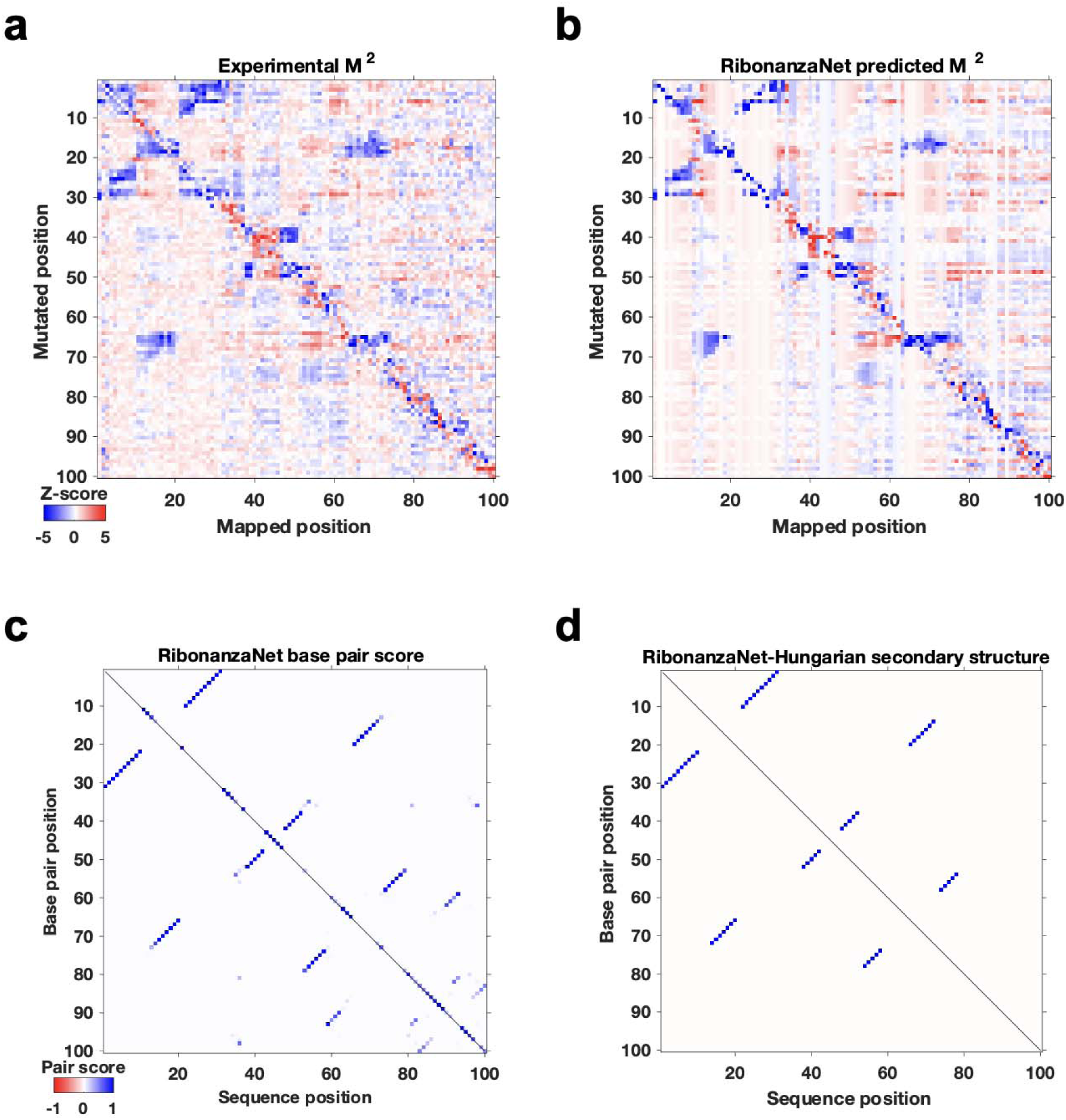
RibonanzaNet predictions for MERS frameshift stimulation element. **(a)** Experimental and **(b)** RibonanzaNet-predicted mutate-and-map measurements for MERS FSE element. **(c)** Pair scores output by RibonanzaNet-SS. **(d)** Final secondary structure output after application of Hungarian algorithm to **(c)**. The estimated accuracy values over the predicted structure and over just the crossed pairs are *eF*_1_ = 0.86 and *eF*_1,crossed pair_ = 0.80, respectively.

**Extended Data Figure 11.**
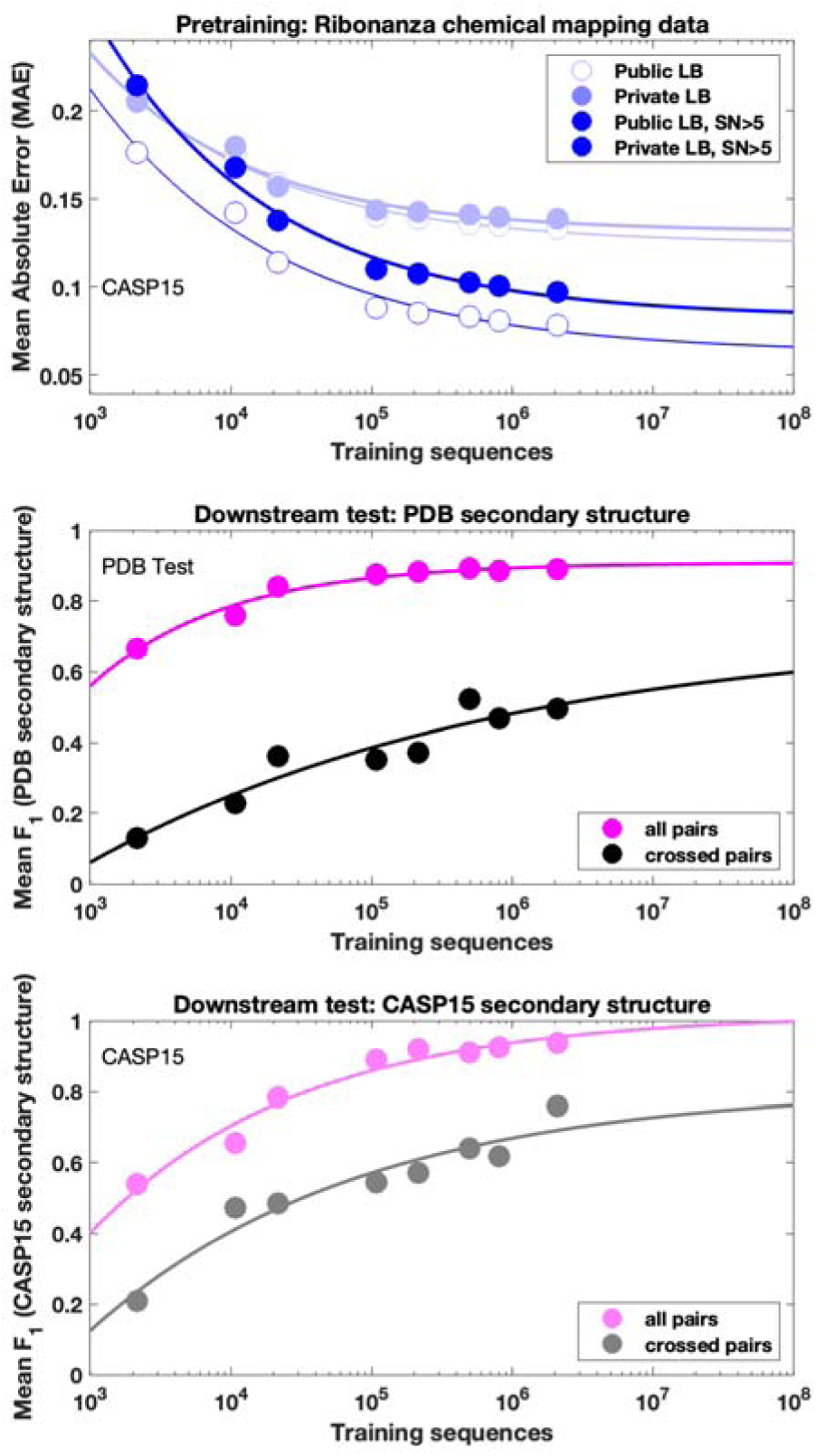
Data ablation studies evaluated on the downstream task of RNA secondary structure. RibonanzaNet was re-trained with randomly sampled subsets of the Kaggle Ribonanza training data (214,831 sequences with either 2A3/DMS profile with signal-to-noise > 1.0; five left most points); a post-competition Ribonanza+ data set for which higher signal-to-noise chemical mapping data were collected (494,111 sequences with either 2A3/DMS profile signal-to-noise > 1.0; sixth point); and the Ribonanza training data supplemented by pseudolabels for the training set (563k noisy training sequences with signal to noise < 1 in both 2A3 and DMS profiles; seventh point) and for the training and test set (2.1M sequences, the final RibonanzaNet model; eighth point). Each of these models was then fine-tuned and evaluated on the same training/test sets as RibonanzaNet-SS, as described in the main text. Chemical mapping MAE scores (light blue, top panel) and *F*_1_ scores for the entire secondary structure (magenta curves, middle and bottom panels) appear to saturate, reflecting intrinsic bounds in both metrics due to data quality and maximal accuracy of 1.0, respectively. However, the MAE computed over higher quality data (signal-to-noise > 5.0, blue, top panel) and the *F*_1, crossed-pair_ scores (black and gray curves, middle and bottom panels) continue to improve with number of training sequences, suggesting good alignment of pre-training and downstream tasks.

**Extended Data Figure 12.**
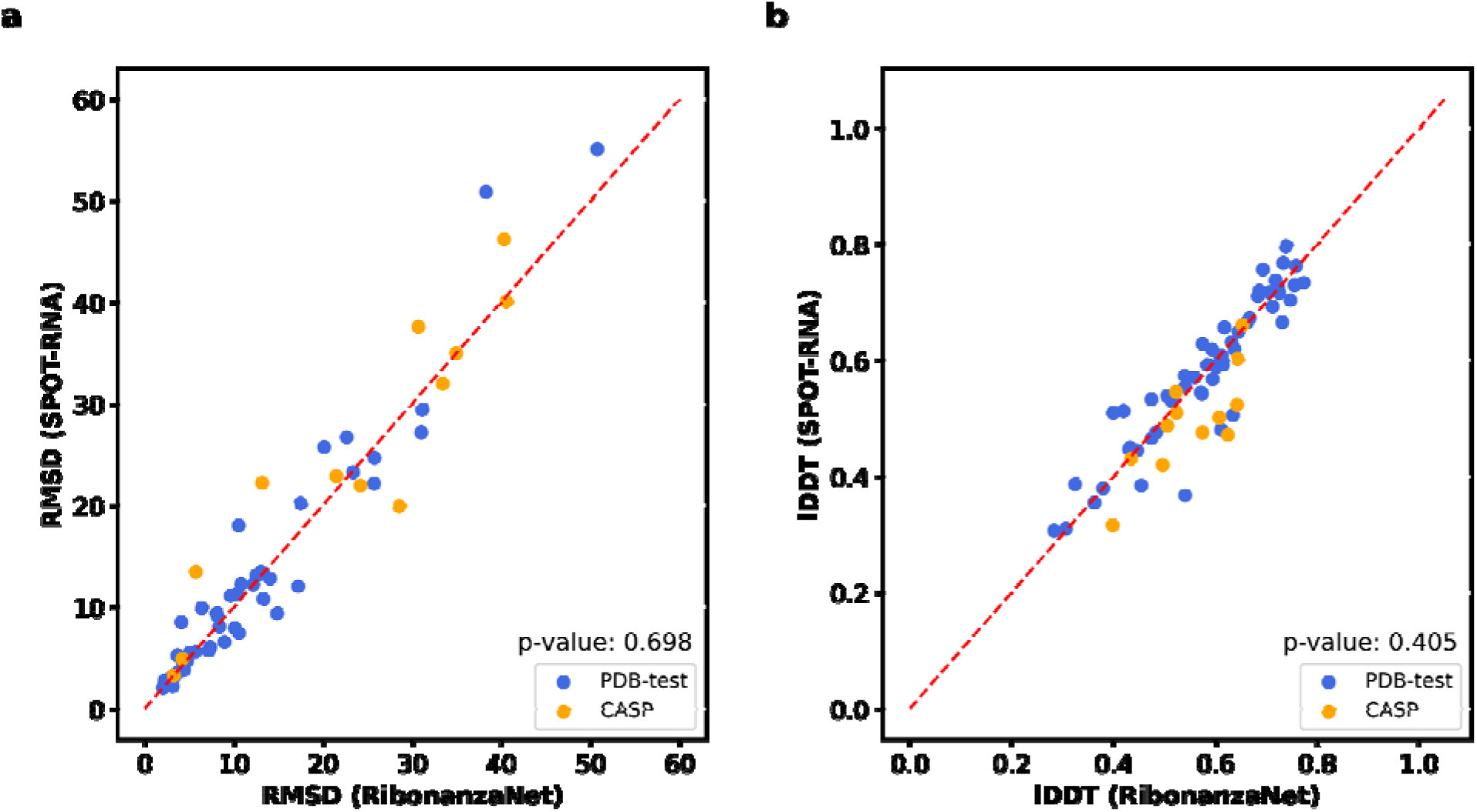
Comparison of accuracy of 3D structures predicted by trRosettaRNA using RibonanzaNet-SS and SPOT-RNA secondary structures. **(a)** RMSD or (**b)** lDDT of structures predicted by trRosettaRNA using secondary structure derived from RibonanzaNet or SPOT-RNA as an input feature. P-values of 0.698 and 0.405 for (a) and (b) are from paired Wilcoxon signed-rank test.

**Extended Data Figure 13.**
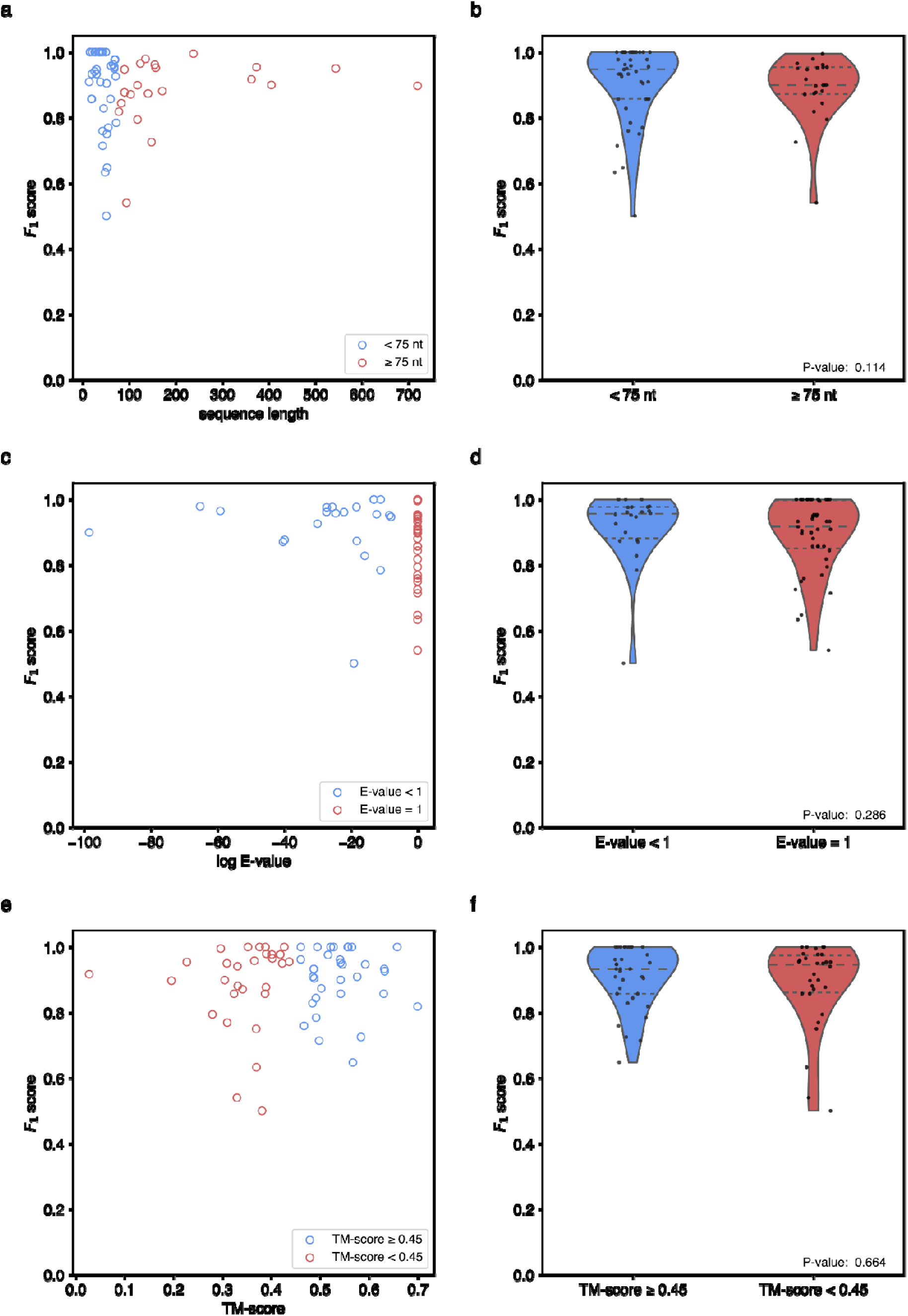
Analysis of RibonanzaNet secondary structure predictions as they relate to sequence and structural parameters. **(a)** Secondary structure *F*_1_ scores for test sequences with respect to the length of the test sequence. **(b)** Comparison of secondary structure *F*_1_ scores for long sequences (greater than or equal to 75 nucleotides) or short sequences (less than 75 nucleotides). (**c)** Secondary structure *F*_1_ test scores with respect to sequence similarity of training sequences. **(d)** Comparison of secondary structure *F*_1_ score values with respect to sequence similarity, separated into sequences with similar sequences in the training dataset (E-value less than 1) and those with no discernible matches by nucleotide BLAST (E-value set to 1). **(e)** Secondary structure *F*_1_ score of test structures with respect to similarity of 3D structures used for fine-tuning, calculated via TM-score with US-align.^55^ **(f)** Comparison of secondary structure *F*_1_ scores with respect to TM-score discretized into test sequences with (TM-score greater than or equal to 0.45) and without (TM-score less than 0.45) a similar 3D structure in the PDB training data utilized during fine tuning. P-values (Wilcoxon rank sum test) for length, E-value and TM-score are 0.114, 0.286, and 0.664 respectively.

## Notes

https://www.kaggle.com/competitions/stanford-ribonanza-rna-folding/data

https://www.kaggle.com/datasets/rhijudas/ribonanza-solutions/

https://www.ncbi.nlm.nih.gov/sra/?term=PRJNA1077399

https://eterna.tech

### Competing Interest Statement

Stanford University is filing patent applications based on concepts described in this paper. R.D. is a cofounder of Inceptive.

### Summary of Updates

(1) Complete description of F1,crossed-pair analysis (pseudoknot prediction accuracy), (2) corrected secondary structure analysis to take into account singlet base pairs, minimal hairpin loop size, and clusters of target sequences, (3) addition of links for ArmNet code & weights, raw data in Sequencing Read Archive, and high-quality Ribonanza+ post-competition data (4) addition of missing ORCIDs for two authors, (5) citation of AlphaFold3 and Rosetta-AllAtom.

